# The NLRP1 and CARD8 inflammasomes detect reductive stress

**DOI:** 10.1101/2022.03.22.485209

**Authors:** Qinghui Wang, Jeffrey C. Hsiao, Noah Yardeny, Hsin-Che Huang, Claire M. O’Mara, Elizabeth L. Orth-He, Daniel P. Ball, Daniel A. Bachovchin

**Affiliations:** Chemical Biology Program, Memorial Sloan Kettering Cancer Center, New York, New York 10065, USA; Pharmacology Program of the Weill Cornell Graduate School of Medical Sciences, Memorial Sloan Kettering Cancer Center, New York, New York 10065, USA; Tri-Institutional PhD Program in Chemical Biology, Memorial Sloan Kettering Cancer Center, New York, New York 10065, USA

**Author notes:** Correspondence to Daniel A. Bachovchin.

## Abstract

The danger signals that activate the NLRP1 and CARD8 inflammasomes have not been fully established. We recently discovered that cytosolic peptide accumulation activates these inflammasomes. In addition, we found that the oxidized form of TRX1 binds to and represses NLRP1, suggesting that NLRP1 also detects a lack of reactive oxygen species, or reductive stress. However, no agents that induce reductive stress were known to test this premise. Here, we identify and characterize several radical-trapping antioxidants, including JSH-23, that induce reductive stress. We show that these compounds accelerate the proteasome-mediated degradation of the repressive N-terminal fragments of NLRP1 and CARD8, releasing the inflammasome-forming C-terminal fragments from autoinhibition. Moreover, we found that reductive stress and peptide accumulation together trigger much more intense inflammasome activation than either signal alone. Overall, this work validates chemical probes that induce reductive stress, and establishes reductive stress alongside peptide accumulation as the key inflammasome-activating danger signals.

## INTRODUCTION

NLRP1 (nucleotide-binding domain leucine-rich repeat pyrin domain-containing 1) and CARD8 (caspase activation and recruitment domain-containing 8) are related human germline-encoded pattern-recognition receptors (PPRs) that sense intracellular danger signals, form multiprotein complexes called inflammasomes, and trigger caspase-1-dependent pyroptosis (Broz and Dixit, 2016; Taabazuing et al., 2020). Although recent research has greatly expanded our understanding of these receptors, the identities of the NLRP1- and CARD8-activating danger signals, and whether they are the same or different, have not yet been fully elucidated (Bachovchin, 2021).

NLRP1 and CARD8 have similar C-terminal regions that consist of FIIND (function-to-find) and CARD (caspase activation and recruitment) domains (**Fig. S1A**). The FIINDs undergo autoproteolysis between ZU5 (ZO-1 and UNC5) and UPA (conserved in UNC5, PIDD, and ankyrin) subdomains, creating N terminal (NT) and C terminal (CT) fragments that remain non-covalently associated (D’Osualdo et al., 2011; Finger et al., 2012; Frew et al., 2012). The proteasome-mediated degradation of the NT fragments releases the CT fragments from autoinhibition in a process referred to as “functional degradation” (**Fig. S1B**). Each freed CT fragment is then sequestered in a ternary complex with one copy of the full-length (FL) PRR and one copy of dipeptidyl peptidase 8 or 9 (DPP8/9), likely to prevent inflammasome formation during homeostatic protein turnover (Hollingsworth et al., 2021; Huang et al., 2021; Sharif et al., 2021; Zhong et al., 2018). NLRP1 and CARD8 inflammasome-activating stimuli accelerate the rate of NT degradation and/or destabilize the DPP8/9 ternary complex, thereby releasing a sufficient level of free CT fragments to self-oligomerize and form an inflammasome (**Fig. S1B**).

The primordial functions of NLRP1 and CARD8 have not been definitively established. It is currently thought that they detect structures and/or activities of pathogens (Mitchell et al., 2019). Consistent with this idea, several pathogen-derived stimuli, including pathogen proteases, E3 ligases, and long dsRNA, have been shown to activate NLRP1 and/or CARD8 (Boyden and Dietrich, 2006; Hornung et al., 2009; Robinson et al., 2020; Sandstrom et al., 2019; Tsu et al., 2020; Wang et al., 2021). However, none of these stimuli activate all NLRP1 alleles in humans and rodents (rodents do not have a CARD8 homolog), suggesting that each NLRP1 allele (as well as CARD8) may sense entirely distinct pathogen-associated features. Alternatively, it is possible that NLRP1 and CARD8 proteins sense a specific perturbation of a fundamental, but as yet unknown aspect of cellular homeostasis (Bachovchin, 2021). Supporting this idea, potent DPP8/9 inhibitors, including Val-boroPro (VbP), activate all functional NLRP1 and CARD8 proteins in humans and rodents by both accelerating the rate of NT degradation and destabilizing the DPP8/9 ternary complexes (**Fig. S1B**) (Chui et al., 2019; Gai et al., 2019; Hollingsworth et al., 2021; Huang et al., 2021; Johnson et al., 2018; Okondo et al., 2017; Sharif et al., 2021; Zhong et al., 2018). DPP8/9 are serine proteases that cleave XP dipeptides (X is any amino acid) from the N-termini of polypeptides (Geiss-Friedlander et al., 2009; Griswold et al., 2019; Tang et al., 2009). Intriguingly, XP peptides, which weakly bind to and inhibit DPP8/9, alone can activate the CARD8 inflammasome, suggesting they may be an endogenous danger-associated signal (Rao et al., 2022). Based on these findings, we have hypothesized that NLRP1 and CARD8 evolved to sense a pathogen-induced “damage state” that is intimately related to DPP8/9 and XP peptides (Bachovchin, 2021; Rao et al., 2022).

We recently discovered that stresses in addition to DPP8/9 inhibition likely contribute to this damage state as well. For example, we found that the non-selective metallo-aminopeptidase (AP) inhibitor bestatin methyl ester (MeBs), which induces the accumulation of many peptides (but not those with XP N-termini), accelerates the rate of NT degradation and thereby synergizes with VbP to induce substantially more pyroptosis (**Fig. S1B**) (Chui et al., 2019; Orth-He et al., 2022). This work indicates that peptide build-up in general (i.e., not just XP peptide build-up) is an inflammasome-activating danger signal. Moreover, we recently uncovered evidence that NLRP1 also senses a lack of reactive oxygen species (ROS), or reductive stress (**Fig. S1B**). Briefly, NLRP1 has NT nucleotide-binding (NACHT) and leucine-rich (LRR) domains (**Fig. S1**), and these domains associate with the oxidized form of thioredoxin-1 (TRX1) (Ball et al., 2021). We found that genetic loss of TRX1 sensitizes cells to VbP-induced pyroptosis, showing that oxidized TRX1 restrains NLRP1 activation. These results predicted that agents that deplete ROS and reduce TRX1 would synergize with VbP to induce more NLRP1-dependent pyroptosis. However, inducers of such reductive stress have not yet been identified and characterized. It should be noted that CARD8 does not have NT NACHT-LRR domains and does not bind oxidized TRX1, and it is unclear if it also detects reductive stress. Moreover, our model predicts that reductive stress and peptide accumulation both contribute to the same damage state (**Fig. S1B**), but it is not obvious how these signals are related to each other.

Here, we wanted to identify small molecule inducers of reductive stress and evaluate their impact on NLRP1 and CARD8 inflammasome activation. We discovered a panel of related radical-trapping antioxidants, and in particular JSH-23, that induce reductive stress, accelerate NT degradation of both NLRP1 and CARD8, and synergize with VbP to induce more pyroptosis. Moreover, we found that reductive stress and peptide accumulation together strongly activate both NLRP1 and CARD8 in the absence of VbP. Overall, this work not only establishes chemical probes that induce intracellular reductive stress, but also reveals that both NLRP1 and CARD8 detect reductive stress.

## RESULTS

### Ferroptosis inhibitors synergize with VbP

We first wanted to assess the impact of antioxidants on NLRP1 and CARD8 inflammasome activation. We initially tested varying doses of a small panel of commonly used antioxidants on VbP-induced pyroptosis in mouse RAW 264.7 macrophages (NLRP1B-dependent) and in human THP-1 and MV4;11 cell lines (CARD8-dependent) using Cell-Titer Glo (CTG, which measures ATP levels as a proxy for cell viability) (Johnson et al., 2018; Okondo et al., 2018). MeBs, which synergizes with VbP to induce more pyroptosis in these cell lines, was used as a positive control in this experiment (Chui et al., 2019; Orth-He et al., 2022). Consistent with our previous results (Ball et al., 2021), we found that dithiothreitol (DTT), N-acetyl cysteine (NAC), reduced L-glutathione (GSH), and trolox did not impact VbP-induced NLRP1 or CARD8 activation (**Fig. 1A,B, Fig. S2A**). Intriguingly, we observed that ferrostatin-1 (Fer-1), a lipophilic antioxidant that blocks an iron-dependent form of cell death called ferroptosis (Dixon et al., 2012), unlike other commonly used antioxidants, appeared to slightly synergize with VbP in RAW 264.7 and MV4;11 cells at a high (40 ìM) concentration (**Fig. 1A,B, Fig. S2A**). Consistent with these results, we found that Fer-1, like MeBs and unlike NAC, also increased VbP-induced lactate dehydrogenase (LDH) release and gasdermin D (GSDMD) cleavage, two hallmarks of pyroptosis, in these cells (**Fig. 1C, Fig. S2B**). These data suggested that Fer-1, unlike other known antioxidants, might synergize with VbP, although this synergistic activity was modest and only observed at high doses.

**Figure 1.**
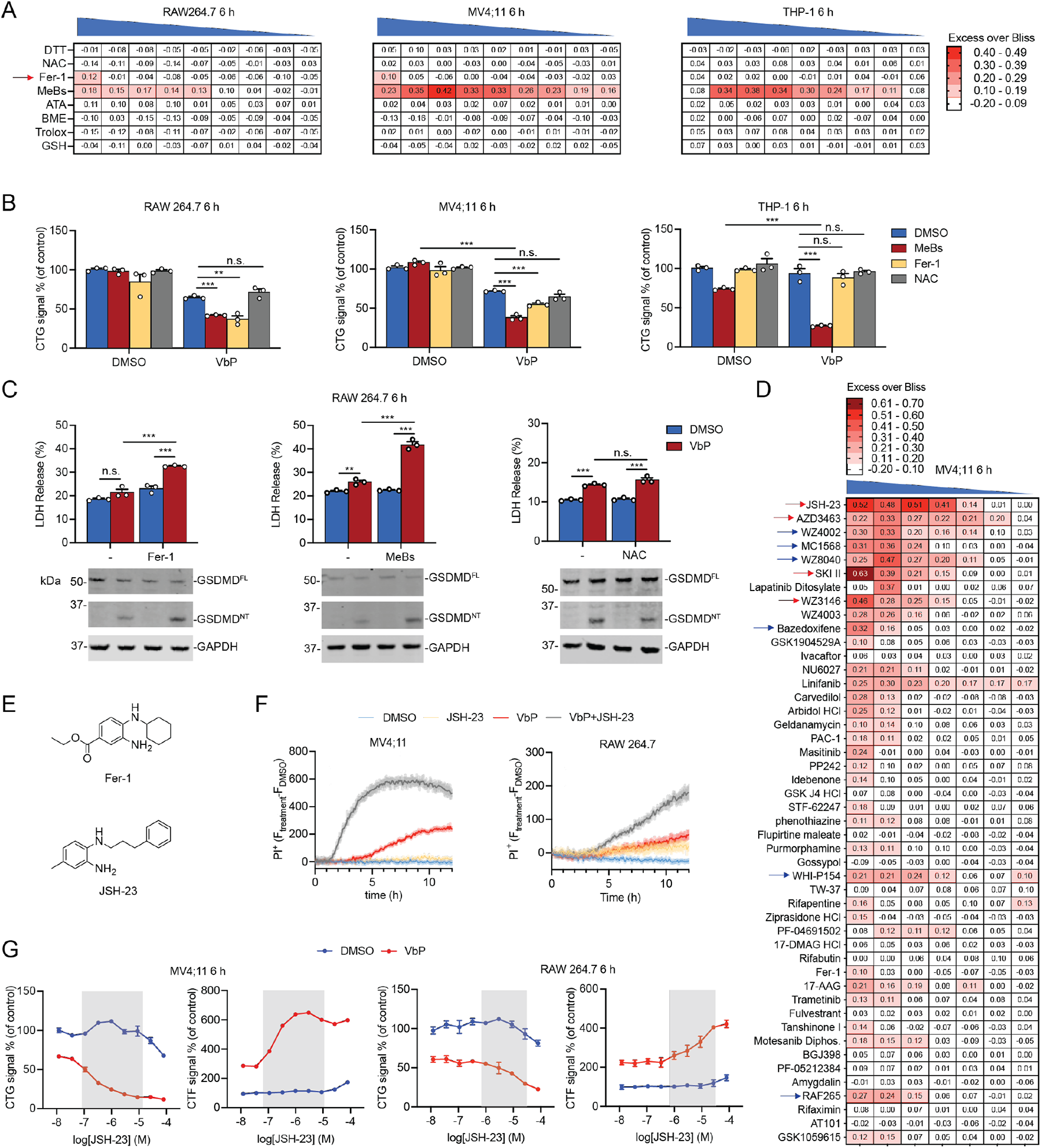
Ferroptosis inhibitors synergize with VbP. (**A**) The indicated cells were treated with the varying doses of the indicated compounds ± VbP (10 μM) for 6 h before cell viability was evaluated by CTG. For each pair of concentrations, we subtracted the predicted Bliss additive effect from the observed inhibition. Values greater than zero indicate synergy. Compounds were all tested in 3-fold dilution series from the following highest doses: DTT (2 mM), NAC (4 mM), Fer-1 (40 μM), MeBs (20 μM), α-Tocopherol acetate (ATA) (2 mM), β-mercaptoethanol (BME) (4 mM), Trolox (400 μM), and GSH (1 mM). (**B**) The indicated cells were treated with MeBs (6.7 μM), Fer-1 (40 μM), and NAC (4 mM) ± VbP (10 μM) for 6 h before CTG analysis. (**C**) RAW 264.7 cells were treated with Fer-1 (40 μM), MeBs (20 μM), and NAC (2 mM) ± VbP (10 μM) for 6 h before cell death was assessed by LDH release and immunoblot analyses. (**D**) MV4;11 cells were treated with varying doses of ferroptosis inhibitors ± VbP (10 μM) for 6h before cell viability was measured by CTG. For each pair of concentrations, we subtracted the predicted Bliss additive effect from the observed inhibition. All compounds were tested in 3-fold dilution series. The highest dose of phenothiazine was 400 μM; the highest doses of MC1568, Amygdalin, Gossypol, Idebenone, SKI II, Flupirtine maleate, Carvedilol, Rifaximin, Masitinib, Ferrostatin-1, WHI-P154, AT101, STF-62247, WZ4002, JSH-23, 17-AAG, NU6027, Motesanib Diphos., Arbidol HCl, PAC-1, GSK J4 HCl, and PP242 were 80 μM; the highest doses of all others were 40 μM. Red and blue arrows indicate compounds further investigated in this manuscript. (**E**) Structures of Fer-1 and JSH-23. (**F**) MV4;11 and RAW 264.7 cells were treated with JSH-23 (10 μM in MV4;11 cells; 1 μM in RAW 264.7 cells), VbP (10 μM), or both before cell death was assessed by PI uptake. (**G**) MV4;11 and RAW 264.7 were treated with the indicated concentrations of JSH-23 and VbP (10 μM) for 6 h before CTG or CTF analyses. Synergistic cell death is highlighted in gray. Data are means ± SEM of 3 replicates. *** *p <* 0.001, ** *p <* 0.01, by two-sided Students *t*-test. n.s., not significant. All data, including immunoblots, are representative of three or more independent experiments. See also Figure S2.

We reasoned that other ferroptosis inhibitors might synergize with VbP more potently than Fer-1. Fortunately, a recent study identified dozens of ferroptosis-suppressing small molecules (Conlon et al., 2021), and we next tested these compounds for their impact on VbP-induced pyroptosis. Notably, we identified several, including JSH-23, AZD3463, SKI II, and WZ3146 (**Fig. 1D**,**E**, **Fig. S2C**), that dramatically synergized with VbP to induce more cell death in MV4;11 cells as measured by CTG. Of these, JSH-23 appeared to have the most synergistic activity. We confirmed that JSH-23 was non-toxic on its own up to 10 ìM after 6 h in both MV4;11 and RAW 264.7 cells, but substantially increased VbP-induced death in both cell types as measured by propidium iodide (PI) uptake (**Fig. 1F**), CTG, and CytoTox-Fluor (CTF, which measures extracellular protease activity after membrane permeabilization) assays (**Fig. 1G**). Similar results were observed with SKI II, WZ3146, and AZD3463 in RAW 264.7 cells as well (**Fig. S2D**). Notably, we found that JSH-23, unlike Fer-1 (**Fig. 1A,B**), induced synergistic cell death with VbP in THP-1 cells (**Fig. S2E,F**). Collectively, these data strongly suggest that several ferroptosis inhibitors, and in particular JSH-23, profoundly enhance VbP-induced cell death.

We next wanted to confirm that these ferroptosis-suppressing small molecules were indeed inducing more pyroptosis and not some other form of cell death. Consistent with amplified pyroptotic cell death, we found that JSH-23, SKI II, AZD3463, WZ3146, and Fer-1 all increased VbP-induced LDH release and GSDMD cleavage in MV4;11 and RAW 264.7 cells (**Fig. 2A,B**). Moreover, JSH-23 induced more LDH release and GSDMD cleavage with VbP in primary resting T cells, which have a functional CARD8 inflammasome (**Fig. S2G**). It should be noted that cleavage of the inflammatory cytokines interleukin-1β and -18 (IL-1β/18), like LDH release and GSDMD cleavage, is typically considered to be a hallmark of pyroptotic cell death. However, inflammasomes that do not use the ASC (apoptosis-associated speck-like protein containing a CARD) adapter protein to bridge to CASP1, which include the CARD8 inflammasome and the NLRP1B inflammasome in RAW 264.7 cells (these cells lack ASC), do not appreciably cleave these cytokines (Ball et al., 2020). Thus, we did not evaluate cytokine cleavage in MV4;11, RAW 264.7, and resting T cells. In contrast to these cells, VbP activates the ASC-dependent human NLRP1 inflammasome in N/TERT-1 keratinocytes, which results in inflammatory cytokine cleavage (Zhong et al., 2018). We found that JSH-23 increased VbP-induced LDH release as well as GSDMD, IL-1β, and IL-18 cleavage in N/TERT-1 keratinocytes (**Fig. 2C**), further indicating that JSH-23 is indeed triggering synergistic pyroptosis.

**Figure 2.**
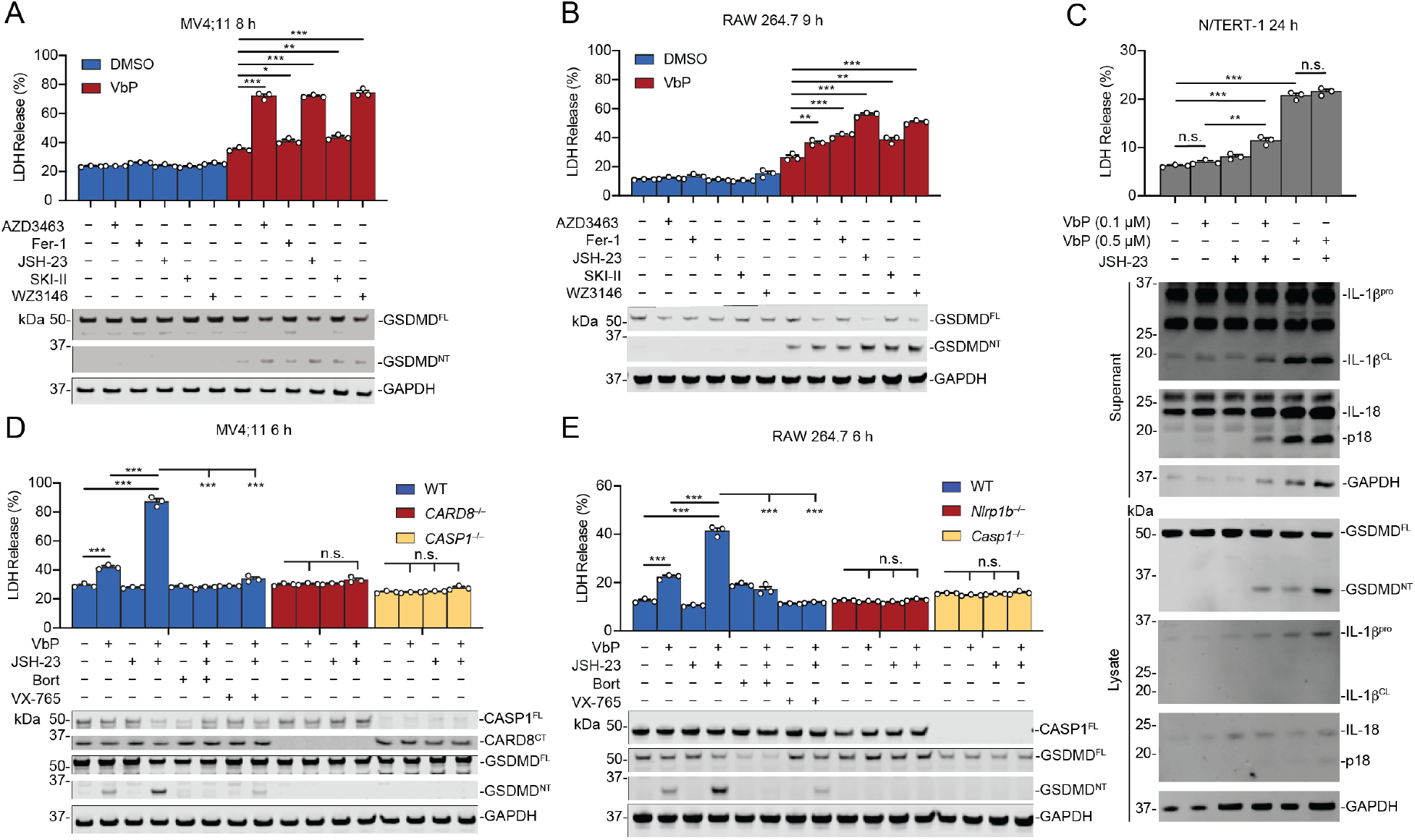
Ferroptosis inhibitors increase CARD8 and NLRP1 inflammasome activation. (**A** and **B**) MV4;11 and RAW 264.7 cells were treated with AZD3463 (1 μM), Fer-1 (20 μM), JSH-23 (1 μM), SKI II (20 μM for MV4;11 cells; 10 μM for RAW 264.7 cells), and WZ3146 (4 μM) ± VbP (10 μM) for the indicated times, followed by LDH release and immunoblot analyses. (**C**) N/TERT-1 keratinocytes were treated with JSH-23 (2 μM) ± VbP for 24 h before LDH release and immunoblot analyses. (**D** and **E**) MV4;11 and RAW 264.7 cells were treated with bortezomib (Bort, 1 μM) or VX-765 (10 μM) for 15 min followed by treatment with VbP (10 μM), JSH-23 (1 μM), or both before LDH release and immunoblot analyses. Data are means ± SEM of 3 replicates. *** *p* < 0.001, ** *p* < 0.01, * *p* < 0.05 by two-sided Students *t*-test. n.s., not significant. All data, including immunoblots, are representative of three or more independent experiments. See also Figure S3.

We next sought to show that this pyroptotic cell death was due to the activation of NLRP1 and CARD8 inflammasomes. Indeed, we found that *CARD8^−/−^* and *CASP1^−/−^* MV4;11 (**Fig. 2D**, **Fig. S3A**,**B**), *CARD8^−/−^* and *CASP1^−/−^* THP-1 cells (**Fig. S3C**,**D**), *Nlrp1b^−/−^* and *Casp1^−/−^* RAW 264.7 cells (**Fig. 2E**, **Fig. S3E**), and *NLRP1^−/−^* N/TERT-1 keratinocytes (**Fig. S3F**,**G**) were completely resistant to the combination of VbP and these ferroptosis inhibitors. The proteasome inhibitor bortezomib and caspase-1 inhibitor VX-765 block NLRP1- and CARD8-dependent cell death by inhibiting NT fragment degradation and GSDMD cleavage, respectively. Again consistent with NLRP1 and CARD8 activation, both bortezomib and VX-765 blocked VbP plus JSH-23-induced cell death in MV4;11, RAW 264.7, and THP-1 cells (**Fig. 2D,E**, **Fig. S3D**). Thus, these ferroptosis-suppressing small molecules synergize with VbP to trigger more robust activation of the NLRP1 and CARD8 inflammasomes.

We wanted to determine if JSH-23 synergy was specific to VbP-induced NLRP1 and CARD8 activation, or it increased pyroptosis induced by other inflammasome activators. Notably, anthrax lethal toxin (LT) directly cleaves the mouse NLRP1B (allele 1) protein, inducing the N-end rule mediated degradation of the NT fragment. We found that JSH-23, if anything, slightly suppressed LT-induced pyroptosis (**Fig. S3H**). In addition, we found that JSH-23 had no impact on lipopolysaccharide (LPS) plus nigericin or imiquimod-induced NLRP3 activation in THP-1 cells (**Fig. S3I**), nor on LPS plus nigericin-induced NLRP3 activation or flagellin-induced NAIP/NLRC4 activation in RAW 264.7 cells stably expressing ASC (RAW 264.7^ASC^) (**Fig. S3J**). Thus, JSH-23 does not generally lead to greater inflammasome activation, but its effect appears specific to the VbP-induced pathway.

### JSH-23 accelerates NT degradation

We next wanted to investigate if JSH-23 increased inflammasome activation by either accelerating NT degradation or destabilizing the DPP8/9 ternary complexes (**Fig. S1B**). We previously generated *DPP8/9^−/−^* THP-1 cells, which lack these repressive complexes and are completely resistant to VbP (Okondo et al., 2017). We found that JSH-23 induced additional lytic cell death and GSDMD cleavage in these cells (**Fig. 3A,B**), which was blocked by the proteasome inhibitor bortezomib (**Fig. S4A**). In addition, we found that JSH-23 treatment of *CASP1^−/−^* MV4;11 cells (knockout cells were used to prevent any pyroptotic cell death) induced a slight reduction in the amount of CARD8^NT^ fragment by immunoblotting (**Fig. 3C,S4B**). It should be noted that only a small amount of the free CARD8^CT^ is required to induce pyroptosis (Chui et al., 2020; Chui et al., 2019; Sandstrom et al., 2019), and therefore even slight degradation is likely physiologically important. Similarly, we only found that JSH-23 induces slight degradation of NLRP1 in N/TERT-1 keratinocytes as evaluated by immunoblotting (**Fig. S4C**). Regardless, these degradation results, coupled with the proteasome blockade and *DPP8/9* knockout data above, collectively indicate that JSH-23 induces synergistic pyroptosis, at least in part, by accelerating NT degradation.

**Figure 3.**
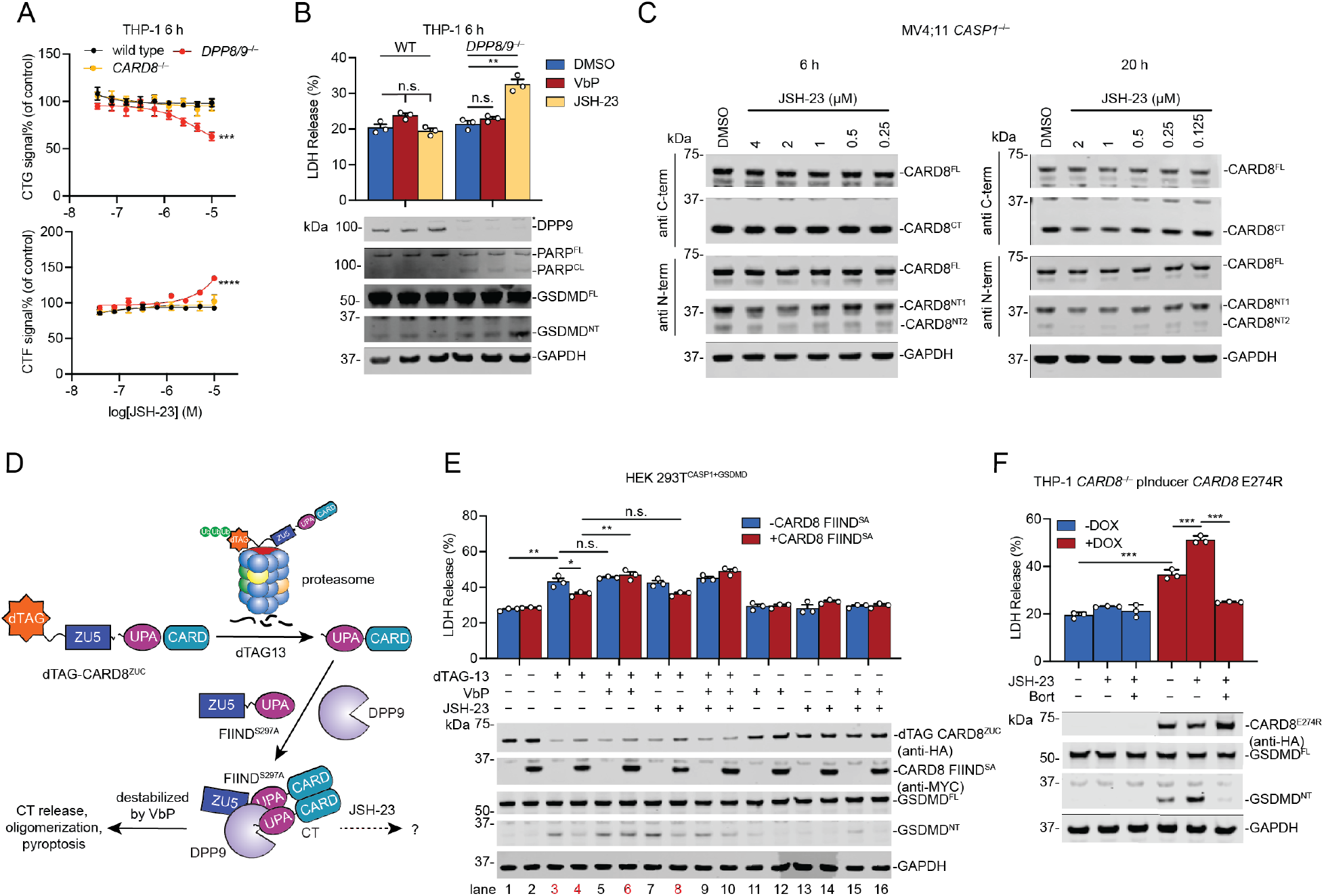
JSH-23 accelerates NT degradation. (**A** and **B**) WT, *DPP8/9^−/−^*, or *CARD8^−/−^* THP-1 cells were treated with VbP (10 μM) or JSH-23 (2 μM in **B**) for 6 h prior to CTG (**A**) and LDH release and immunoblot analyses (**B**). (**C**) *CASP1^−/−^* MV4;11 cells were treated with the indicated concentrations of JSH-23 for 6 h (left) or 20 h (right). CARD8 protein levels were then evaluated by immunoblotting. CARD8 NT1 and NT2 are from different splice isoforms of CARD8. (**D**) Schematic of dTAG-based assay used to evaluate the ability of compounds to destabilize the CARD8-DPP8/9 ternary complex in cells. (**E**) HEK 293T cells stably expressing CASP1 and GSDMD were transiently transfected with plasmids encoding dTAG-CARD8^ZUC^ and CARD8 FIIND^S297A^. Cells were treated with dTAG-13 (500 nM), VbP (10 μM), and JSH-23 (2 μM) for 3 h before LDH release and immunoblot analyses. (**F**) *CARD8^−/−^* THP-1 cells containing a dox-inducible CARD8 E274R mutant that does not bind to DPP8/9 were pre-treated with or without DOX (100 ng/mL, 16 h), and then treated with JSH-23 (5 μM) ± Bort (1 μM) for 6 h, followed by LDH release and immunoblot analyses. Data are means ± SEM of 3 replicates. *** *p* < 0.001, ** *p* < 0.01, * *p* < 0.05 by two-sided Students *t*-test. n.s., not significant. All data, including immunoblots, are representative of three or more independent experiments. See also Figure S4.

Even so, we reasoned that it remained possible that JSH-23 might also destabilize the DPP8/9 ternary complex in wild-type cells. We therefore next tested JSH-23 in an assay we previously developed to assess ternary complex disruption in cells (**Fig. 3D**) (Sharif et al., 2021). This assay relies on the degron tag system, in which the small molecule dTAG-13 induces the rapid degradation of proteins fused to degron tags (dTAGs) (Nabet et al., 2018). Here, we appended a dTAG to the N-terminus of CARD8 <u>Z</u>U5-<u>U</u>PA-<u>C</u>ARD region to generate a dTAG-13-activatable fusion protein (dTAG-CARD8^ZUC^, **Fig. 3D**), and transiently expressed this fusion protein in HEK 293T cells stably expressing GSDMD and CASP1 (HEK 293T^CASP1+GSDMD^). Notably, dTAG-CARD8^ZUC^ is insensitive to VbP because it lacks the N-terminal disordered region required for DPP8/9 inhibitor-induced NT degradation (Chui et al., 2020; Sharif et al., 2021). As expected, dTAG-13, but not VbP, JSH-23, or VbP plus JSH-23, treatment triggered dTAG-CARD8^ZUC^ degradation and induced pyroptosis in these cells (**Fig. 3E**, lane 3). Importantly, co-expression of an autoproteolysis-defective S297A mutant CARD8 FIIND domain (FIIND^SA^) blocked this pyroptosis by sequestering the dTAG13-generated free CARD8^CT^ in a repressive ternary complex (**Fig. 3D**, **Fig. 3E**, lane 4 versus 3). We found that VbP destabilized this repressive complex and restored LDH release and GSDMD cleavage (**Fig. 3E**, lane 6 versus 4), but JSH-23 had no impact on this assay (**Fig. 3E**, lane 8 versus 4). Thus, JSH-23 does not destabilize the DPP8/9 ternary complex.

We instead predicted that JSH-23 acts solely by accelerating NT fragment degradation, even if it was barely observable by immunoblotting (**Fig. 3C, Fig. S4C**). Moreover, we speculated that JSH-23 does not induce pyroptosis on its own (in the absence of VbP) because the released CT fragments are effectively quenched by the DPP8/9 ternary complex (**Fig. S4D**). To test this idea, we next introduced a construct encoding a doxycycline (DOX)-inducible E274R mutant CARD8 protein into *CARD8^−/−^* THP-1 cells. Importantly, the full-length CARD8 E274R protein cannot bind to DPP8/9, and therefore any released free CT fragments are not sequestered in the ternary complex, but instead form inflammasomes (Sharif et al., 2021) (**Fig. S4D**). Indeed, DOX treatment induced spontaneous pyroptosis in these cells due to the generation of free CARD8^CT^ fragments during homeostatic protein turnover (**Fig. 3F**). Consistent with our hypothesis, we found that JSH-23 induced additional pyroptosis in CARD8 E274R-expressing cells, and bortezomib completely blocked this cell death. Similarly, we found that JSH-23 induced more pyroptosis in *CARD8^−/−^* THP-1 cells ectopically expressing NLRP1 P1214R, which also has attenuated binding to DPP9 (**Fig. S4E**) (Zhong et al., 2018). These data demonstrate that JSH-23 alone induces a danger signal that accelerates the degradation of the NT fragments, but that the DPP8/9 ternary complex prevents the resulting free CT fragments from inducing pyroptosis. VbP synergizes with JSH-23 because it prevents DPP8/9 from sequestering these newly released CT fragments.

### Synergistic compounds are radical trapping antioxidants

Our next objective was to identify the danger state induced by the ferroptosis inhibitors. Before exploring their antioxidant activities, we first confirmed that JSH-23 and SKI II do not inhibit recombinant DPP9 (**Fig. 4A**), and that JSH-23, SKI II, AZD3463, and WZ3146 do not inhibit DPP8/9 activity in living HEK 293T cells (**Fig. 4B**). 8j is a selective DPP8/9 inhibitor and was used as a control in this experiment (Van Goethem et al., 2008). As mentioned previously, non-selective inhibitors of aminopeptidases, including MeBs, synergize with VbP to induce more NLRP1 and CARD8 inflammasome activation (Chui et al., 2019; Orth-He et al., 2022) We also confirmed that the ferroptosis inhibitors, unlike MeBs, do not block aminopeptidase activity in cells (**Fig. 4C**). Lastly, we showed that these compounds, unlike the proteasome inhibitors bortezomib and MG132, do not impact overall proteasome activity in cells (**Fig. 4D**). Thus, these compounds do not interfere with any of the known pathways that regulate NLRP1 and CARD8 inflammasome activation.

**Figure 4.**
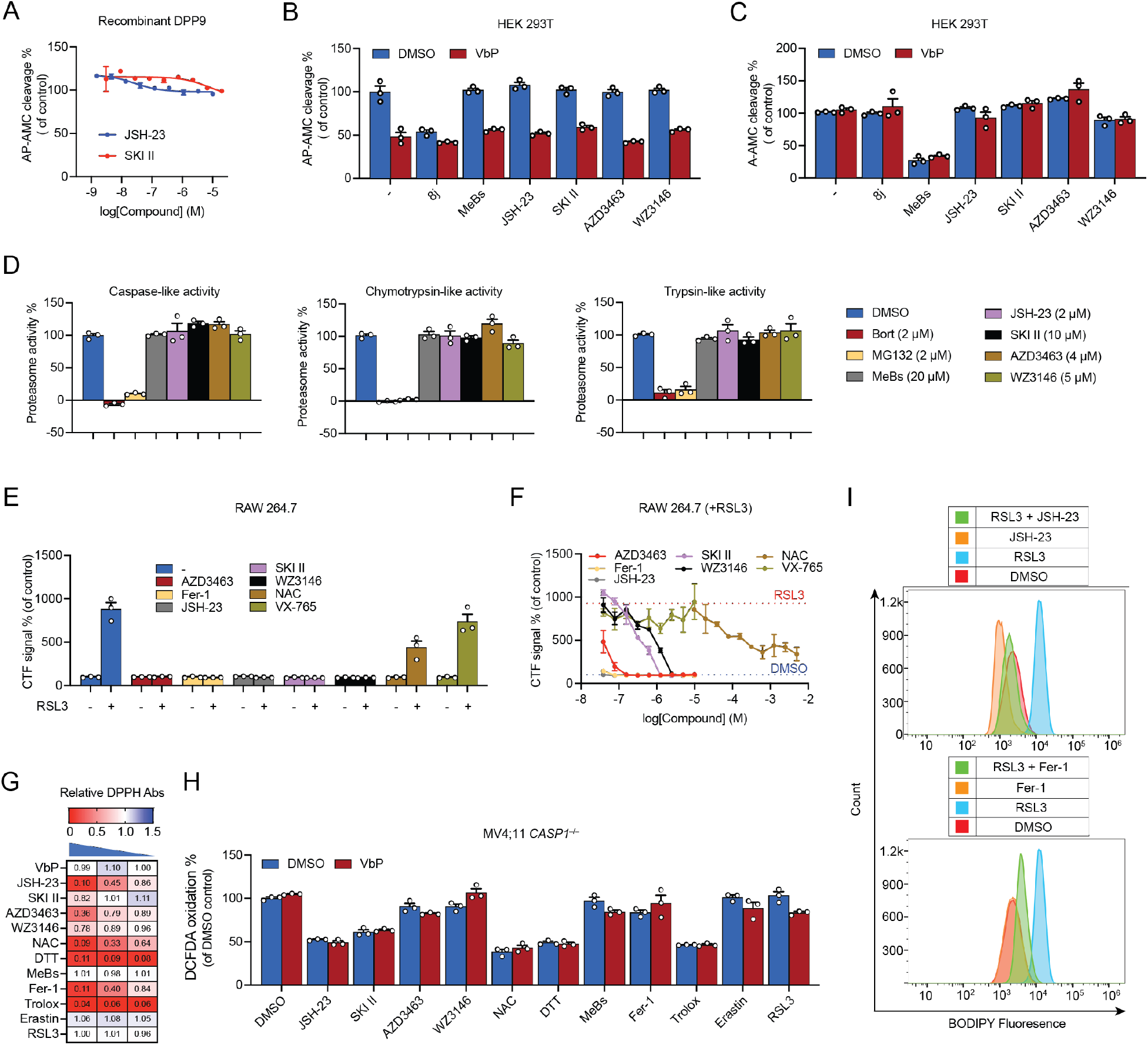
Synergistic compounds are radical trapping agents that induce reductive stress. (**A**) The indicated concentrations of JSH-23 and SKI-II do not inhibit the activity of recombinant DPP9. (**B** and **C**) HEK 293T cells were treated with 8j (10 μM), MeBs (20 μM), JSH-23 (5 μM), SKI II (10 μM), AZD3463 (1 μM), WZ3146 (4 μM) ± VbP (10 μM) before assaying for AP-AMC (**B**) or A-AMC (**C**) cleavage. (**D**) The ferroptosis inhibitors do not impact proteasome activity in the cell-based proteasome-Glo assay. (**E** and **F**) RAW 264.7 cells were treated with the indicated compounds ± RSL3 (0.25 μM). Cell death was evaluated by CTF after 4.5 h. All compounds in **E** were tested at 5 μM (except for NAC at 1.2 mM). (**G**) VbP (10 μM), JSH-23 (10 μM), SKI II (10 μM), AZD3463 (10 μM), WZ3146 (10 μM), NAC (0.5 mM), DTT (0.5 mM), MeBs (10 μM), Fer-1 (25 μM), Trolox (200 μM), Erastin (10 μM), RSL3 (0.25 μM) were tested for radical trapping activity using a cell-free DPPH assay. The highest concentration for each compound is indicated above, which was then diluted 5-fold serially. (**H**) VbP (10 μM), JSH-23 (4 μM), SKI II (20 μM), AZD3463 (4 μM), WZ3146 (0.8 μM), NAC (1 mM), DTT (1 mM), MeBs (20 μM), Fer-1 (0.8 μM), Trolox (400 μM), Erastin (20 μM) and RSL3 (0.5 μM) ± VbP (10 μM) were tested for their impact on the oxidation of the cell-permeable H_2_DCFDA probe in *CASP1^−/−^* MV4;11 cells. (**I**) *Casp1^−/−^* RAW 264.7 cells were treated with JSH-23 (1 μM), Fer-1 (1 μM), RSL3 (1 μM), or the indicated combinations for 2 h prior to assessing the oxidation of the C11 BODIPY 581/591 probe. Data are means ± SEM of 3 replicates. All data are representative of three or more independent experiments. See also Figure S5.

We next sought to verify that these compounds indeed inhibited ferroptosis. The small molecule RSL3 induces ferroptosis by inhibiting glutathione peroxidase 4 (GPX4), thereby causing the accumulation of lipid peroxides and cell death (Yang et al., 2014). We found that all five of the reported ferroptosis inhibitors completely blocked RSL3-induced death in RAW 264.7 cells (**Fig. 4E,F**, **Fig. S5A,B**). In contrast, NAC only partially blocked ferroptosis, even at high (millimolar) concentrations, and the caspase-1 inhibitor VX-765 had no impact in this experiment. Fer-1 and JSH-23 were the most potent ferroptosis inhibitors, blocking cell death at low nanomolar concentrations (< 100 nM), whereas SKI II and WZ3146 required low micromolar concentrations to effectively block cell death. Thus, these compounds all share the ability to inhibit ferroptosis. Notably, their ferroptosis-blocking activities generally correlate with their pyroptosis-inducing activities, with the exception that Fer-1 is far more effective in suppressing ferroptosis.

We next wondered if the pyroptosis-inducing abilities of these compounds might correlate more strongly with their abilities to quench free radicals. As such, we next tested the ability of these compounds to directly scavenge free radicals *in vitro* using the cell-free 2,2-diphenyl-1-picrylhydrazyl (DPPH) assay. JSH-23 and Fer-1, and to a lesser extent AZD3463, SKI II, and WZ3146, indeed scavenged free radicals (**Fig. 4G**). However, we found that NAC, DTT, and Trolox, which do not induce synergistic pyroptosis, also had potent activity in this assay. Thus, the synergistic compounds all scavenge radicals *in vitro*, but this activity alone does not predict their impact on NT degradation in cells. As expected, VbP and MeBs had no impact on DPPH absorbance, consistent with their distinct mechanisms of action (**Fig. 4G**).

We next wanted to assess the impact of these compounds in cell-based antioxidant activity assays. The cell permeable 2’,7’-dichlorodihydrofluorescein diacetate dye (DCFDA), which fluoresces upon oxidation, is commonly used to measure intracellular ROS. We found that JSH-23 and SKI II slowed DCFDA oxidation, but so did NAC, DTT, and Trolox (**Fig. 4H, Fig. S5C**). Moreover, AZD3463 and WZ3146 did not show activity in this assay. Thus, DCFDA oxidation further supports the idea that JSH-23 is an antioxidant, but also that this assay likely does not measure the form of intracellular ROS that suppresses pyroptosis. We next tested the impact of these compounds on the oxidation of the C11 BODIPY probe, which acts as a sensor of lipid peroxidation in cells. We found that JSH-23, AZD3463, SKI II, WZ3146, but not Fer-1, slowed the basal oxidation of this probe in cells (**Fig. 4I, Fig. S5D**). Moreover, we found that these compounds, except WZ3146, substantially suppressed RSL3-induced oxidation of this probe. This data further supports the idea that the synergistic compounds are all antioxidants, and their activities in this assay, as in the ferroptosis blockade assay, generally correlate with their pyroptosis-inducing abilities. Overall, these results confirm that JSH-23 and the other ferroptosis inhibitors act as radical trapping antioxidants (RTAs), even if these commonly used assays do not precisely measure the intracellular ROS that regulates inflammasome activation.

### Structure-activity relationship of the RTAs

JSH-23, SKI II, WZ3146, AZD3463, and Fer-1 all have aromatic secondary amines that can potentially scavenger free radicals (**Fig. S6A**, colored red) (Conlon et al., 2019; Ingold and Pratt; Shah et al.), but are otherwise not structurally related. We noticed that several additional hits from the primary screen similarly had such secondary aromatic amines (WZ4002, WZ8040, WHI-P154, and RAF265), but some other hits did not (MC1568, bazedoxifene) (**Fig. 1D**, blue arrows, **Fig. S6A**). Interestingly, only the compounds with the secondary aromatic amines exhibited synergistic pyroptotic activity in confirmation assays in MV4;11 cells (**Fig. S6B**). To directly investigate the importance of this moiety, we next purchased analogs of JSH-23 itself (**Fig. 5A**). Notably, compound **1**, which has a second phenyl group directly attached to the secondary amine, retained synergistic activity in CTG assays in both MV4;11 and RAW 264.7 cells (**Fig**. **5B**,**C**). However, compound **2**, which is identical to **1** but without secondary amine, lacked all activity. The primary amine was also critical, as diarylamine (compound **7**) was inactive. As expected, compounds **6** and **11**, which have substituents on the secondary amine similar to JSH-23, were also active. Lastly, the electronics of the aromatic ring of **6** appeared to be critical, as replacement of the electron-donating methyl group with an electron-withdrawing cyano group (compound **4**) ablated all activity. We confirmed the CTG results indeed reported pyroptotic cell death, as the active compounds increased VbP-induced PI uptake in MV4;11 cells (**Fig. 5D**) as well as LDH release and GSDMD cleavage in both MV4;11 cells and RAW 264.7 cells (**Fig. 5E, Fig. S6C**). As expected, this cell death was completed blocked by bortezomib and VX-765 (**Fig. S6D**,**E**).

**Figure 5.**
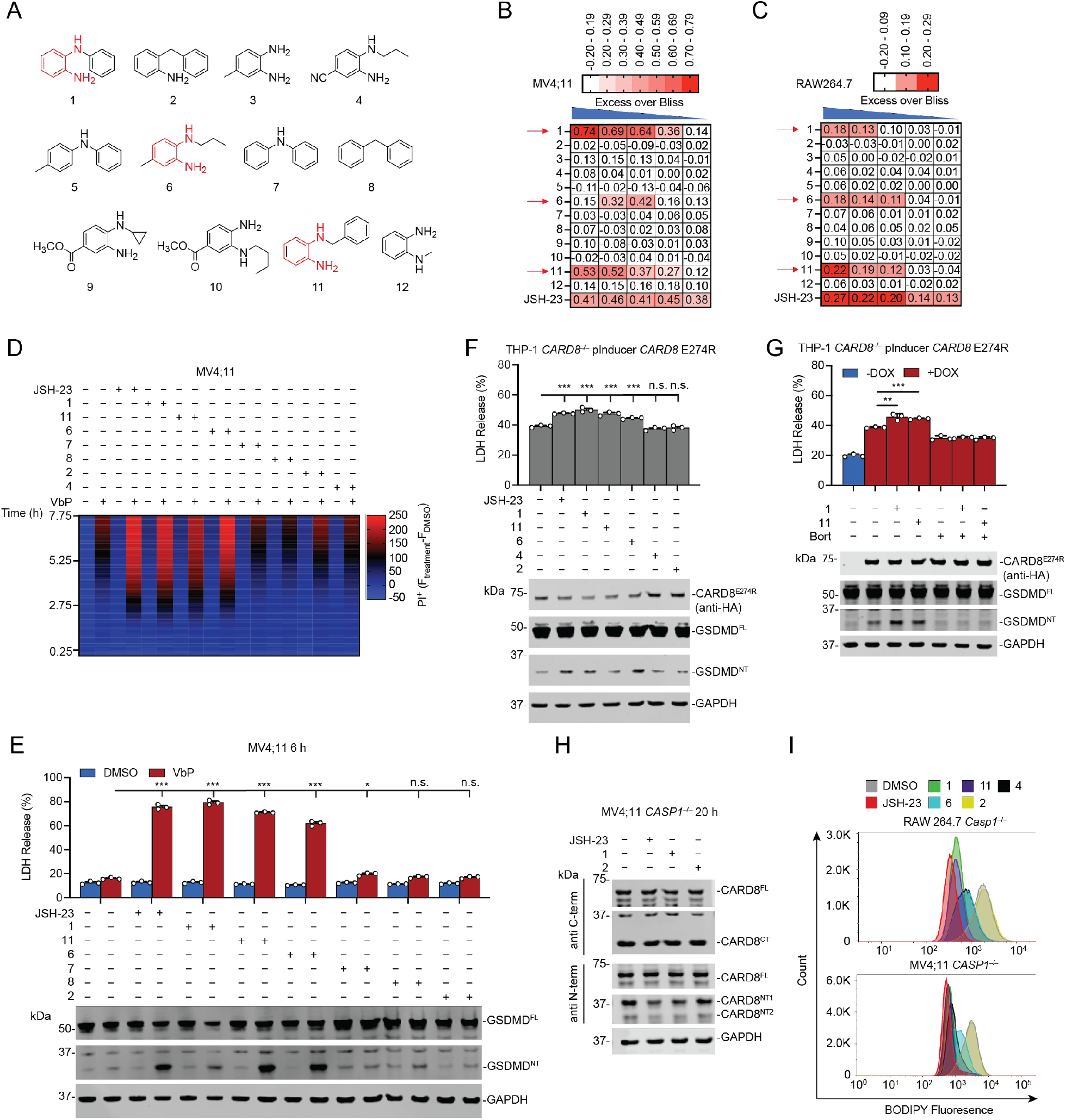
Structure-activity relationship of JSH-23. (**A**) Structures of JSH-23 derivatives. (**B** and **C**) MV4;11 (**B**) and RAW 264.7 (**C**) cells were treated with the varying doses of the indicated compounds (highest concentration for each is 40 μM, followed by serial 3-fold dilutions) ± VbP (10 μM) for 6 h before cell viability was evaluated by CTG. For each pair of concentrations, we subtracted the predicted Bliss additive effect from the observed inhibition. (**D** and **E**) MV4;11 cells were treated with the indicated JSH-23 derivative (5 μM) ± VbP (10 μM) prior to assessing cell death by PI uptake over 8 h (**D**) or by LDH release and immunoblot analyses after 6 h (**E**). (**F** and **G**) *CARD8^−/−^* THP-1 cells containing a DPP9 non-binding CARD8 E274R mutant were treated with DOX (100 ng/mL, 16 h, **F**) or with or without DOX (100 ng/mL, 16 h, **G**) followed by Bort (1 μM), JSH-23 (2 μM), compounds **1**, **2**, **4**, **6**, **11**, (all were tested at 5 μM), or the indicated combinations for 6 h prior to LDH release and immunoblot analyses. (**H**) *CASP1^−/−^* MV4;11 cells were treated with JSH-23, **1**, or **2** (all 2 ìM) for 20 h before immunoblot analysis. (**I**) Compounds **1**, **2**, **4**, **6**, **11** (all at 5 μM for 2 h) were tested for their impact on lipid ROS in *Casp1^−/−^* RAW 264.7 using the C11 BODIPY 581/591 probe. Data are means ± SEM of 3 replicates. *** *p* < 0.001, ** *p* < 0.01, * *p* < 0.05 by two-sided Students *t*-test. n.s., not significant. All data, including immunoblots, are representative of three or more independent experiments. See also Figure S6.

Not surprisingly, the synergistic compounds, but not the inactive compound **2**, appeared to have mechanisms of action like JSH-23. For example, the synergistic compounds induced additional bortezomib-sensitive pyroptosis in *CARD8^−/−^* THP-1 cells ectopically expressing the DPP9-non-binding mutant CARD8 E274R (**Fig. 5F,G**), induced some visible CARD8^NT^ depletion by immunoblotting (**Fig. 5H, Fig. S6F**), trapped free radicals in the DPPH assay (**Fig. S6G**), and acted as antioxidants in the cell-based C11 BODIPY and DCFDA assays (**Fig. 5I, Fig. S6H**). Overall, these data show that the conjugated amines of JSH-23 are critical for its activity, as these groups likely trap specific ROS species inside cells, induce reductive stress, and accelerate NT degradation.

### AP inhibitors and RTAs trigger inflammasome activation

As mentioned above, we recently discovered that the AP inhibitor MeBs synergizes with VbP to induce more NLRP1 and CARD8 inflammasome activation (Chui et al., 2019), similar to the RTAs studied here. However, MeBs does not trap free radicals (**Fig. 4G**), but instead induces the accumulation of intracellular oligopeptides that accelerate NT degradation (Orth-He et al., 2022). Thus, JSH-23 and MeBs have entirely distinct mechanisms of action. Accordingly, we hypothesized that the combination of MeBs and JSH-23 might induce sufficient NT degradation and CT release to overcome the repressive DPP8/9 ternary complex even in the absence of VbP. To test this premise, we next treated MV4;11 cells with JSH-23 and increasing doses of MeBs. Excitingly, we observed that this drug combination stimulated pyroptosis in WT, but not *CASP1^−/−^*, MV4;11 cells (**Fig. 6A,B, Fig. S7A**). As expected, JSH-23 + MeBs induced degradation of the CARD8^NT^ fragment (**Fig. 6C**), although the additional impact of MeBs on JSH-23-induced degradation is difficult to observe by immunoblotting. Nevertheless, this degradation was critical, as JSH-23 + MeBs-triggered pyroptosis was blocked by VX-765 and bortezomib (**Fig. 6D**). We similarly observed that JSH-23 + MeBs induced inflammasome activation in CARD8-dependent THP-1 cells and OCI-AML2 cells (**Fig. 6E, Fig. S7B**). We found that this drug combination did not appear to impact DPP8/9 activity nor destabilize the CARD8-DPP8/9 ternary complex in cells (**Fig. 6F, Fig. S7C**), but did increase the activation of CARD8 E274R higher than either drug alone (**Fig. 6G**). Overall, these results indicate the AP inhibitors and RTAs together induce sufficient NT degradation to activate the CARD8 inflammasome. Even though we do not see evidence of CARD8-DPP8/9 ternary complex destabilization, we should note that it remains possible that this drug combination might also induce the accumulation of some peptides that slightly weaken this checkpoint in a way that is difficult to directly observe (Orth-He et al., 2022).

**Figure 6.**
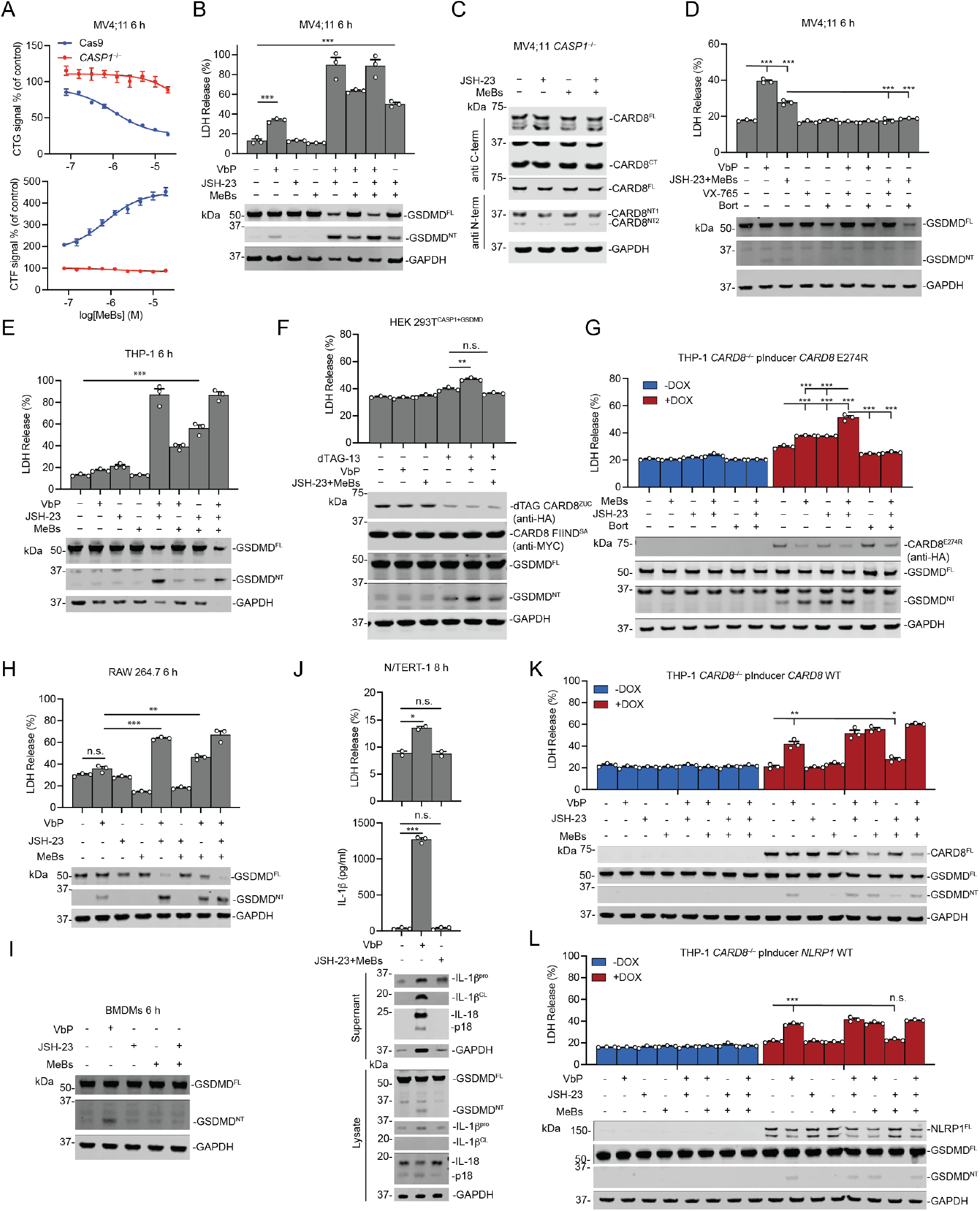
AP inhibitors and RTAs activate CARD8 inflammasome. (**A**) Control (Cas9) or *CASP1^−/−^* MV4;11 cells were treated with JSH-23 (5 μM) and the indicated concentration of MeBs for 6 h before cell viability was evaluated by CTG. (**B**-**E**) The indicated cells were treated with JSH-23 (2 μM), MeBs (10 μM), VX-765 (50 μM), Bort (1 μM), or the specified combinations for 6 h prior to LDH release and immunoblot analyses. VX-765 and Bort were applied 30 min before the inflammasome inducers. (**F**) HEK 293T cells stably expressing CASP1 and GSDMD were transiently transfected with plasmids encoding dTAG-CARD8^ZUC^ and CARD8 FIIND^S297A^. Cells were treated with dTAG-13 (500 nM), VbP (10 μM), JSH-23 (2 μM), and MeBs (10 μM) for 3 h before LDH release and immunoblot analyses. (**G**) *CARD8^−/−^* THP-1 cells containing a DOX-inducible DPP9 non-binding CARD8 E274R protein were treated with or without DOX (100 ng/mL, 16 h) before the addition of JSH-23 (5 μM), MeBs (10 μM), and Bort (1 μM) for 6 h prior to LDH release and immunoblot analyses. Bort was applied 30 min before the inflammasome inducers. (**H**-**J**) The indicated cells were treated with JSH-23 (2 μM), MeBs (10 μM), VbP (10 μM), or the specified combinations for the indicated time intervals before LDH release, IL-1β release, and immunoblotting analysis. (**D** and **E**) *CARD8^−/−^* THP-1 cells containing either a DOX-inducible CARD8 protein or NLRP1 protein were treated with or without DOX (100 ng/mL for CARD8, 1 μg/mL for NLRP1, 16 h) before the addition of JSH-23 (2 μM), MeBs (10 μM), or the specified combinations for 6 h prior to LDH release and immunoblot analyses. Data are means ± SEM of 3 replicates. *** *p* < 0.001, ** *p* < 0.01, * *p* < 0.05 by two-sided Students *t*-test. n.s., not significant. All data, including immunoblots, are representative of three or more independent experiments. See also Figure S7.

In contrast to CARD8, we found that MeBs + JSH-23 did not activate mouse NLRP1B (allele 1) in RAW 264.7 cells (**Fig. 6H**), mouse NLRP1A and/or NLRP1B (allele 2) in C57BL/6 bone marrow-derived macrophages (BMDMs, **Fig. 6I**), or the human NLRP1 inflammasome N/TERT-1 keratinocytes (**Fig. 6J**). To confirm that the inability of JSH-23 + MeBs to activate NLRP1 was due to NLRP1 itself and not the distinct cell types, we next ectopically expressed CARD8 or NLRP1 in *CARD8^−/−^* THP-1 cells. As expected, VbP alone or in combination with MeBs or JSH-23 induced GSDMD cleavage in both cell types, but JSH-23 + MeBs only triggered GSDMD cleavage (albeit modestly) in cells expressing CARD8 (**Fig. 6K**,**L**). Similarly, compounds **1**, **6**, and **11** in combination with MeBs also induced CARD8 inflammasome activation in MV4;11 cells, but not NLRP1B inflammasome activation in RAW 264.7 cells (**Fig. S7D-F**). Collectively, these data indicate that these combinations do not induce sufficient NT degradation and/or ternary complex destabilization to activate NLRP1. These results are consistent with the idea that NLRP1 has a higher activation threshold than CARD8, as discussed below (Hollingsworth et al., 2021; Rao et al., 2022; Sharif et al., 2021).

### XP peptides and RTAs activate NLRP1 and CARD8

We recently described a small molecule called CQ31 that inhibits the aminopeptidases PEPD and XPNPEP1 and thereby causes the intracellular accumulation of XP peptides. Interestingly, CQ31 alone, like the MeBs + JSH-23 combination, activates the human CARD8, but not the human NLRP1, inflammasome (Rao et al., 2022). NLRP1, unlike CARD8, directly interacts with the DPP8/9 active site in the ternary complex (Hollingsworth et al., 2021; Sharif et al., 2021), and we hypothesized that the CQ31-accumulated peptides did not sufficiently destabilize the NLRP1-DPP9 interaction to trigger inflammasome activation, at least in the absence of a stimulus that potently accelerates NT degradation.

We next wondered if the combination of CQ31 and JSH-23 together would induce pyroptosis. As expected, we found JSH-23 synergized with CQ31 to induce more CARD8-dependent pyroptosis in MV4;11 cells (**Fig. 7A**). Notably, CQ31 + JSH-23 induced more cell death than VbP alone, CQ31 alone, or CQ31 + MeBs at this time point. Intriguingly, we found that the combination of JSH-23 + CQ31 also slightly activated the human NLRP1 inflammasome in N/TERT-1 keratinocytes, and the triple combination of JSH-23 + MeBs + CQ31 caused substantial NLRP1 activation (**Fig. 7B, Fig. S7G).** However, this triple drug cocktail was still unable to activate the mouse NLRP1B inflammasome in RAW 264.7 cells (**Fig. S7H**), consistent with previous data that shows the mouse NLRP1 inflammasome has an especially high activation threshold (Cirelli et al., 2014; Ewald et al., 2014; Gai et al., 2019). Regardless, these data collectively indicate that human CARD8 and NLRP1 are activated by the same danger signals, but NLRP1 has a higher activation threshold that requires the simultaneous occurrence of several key inputs (**Fig. 7C,D**).

**Figure 7.**
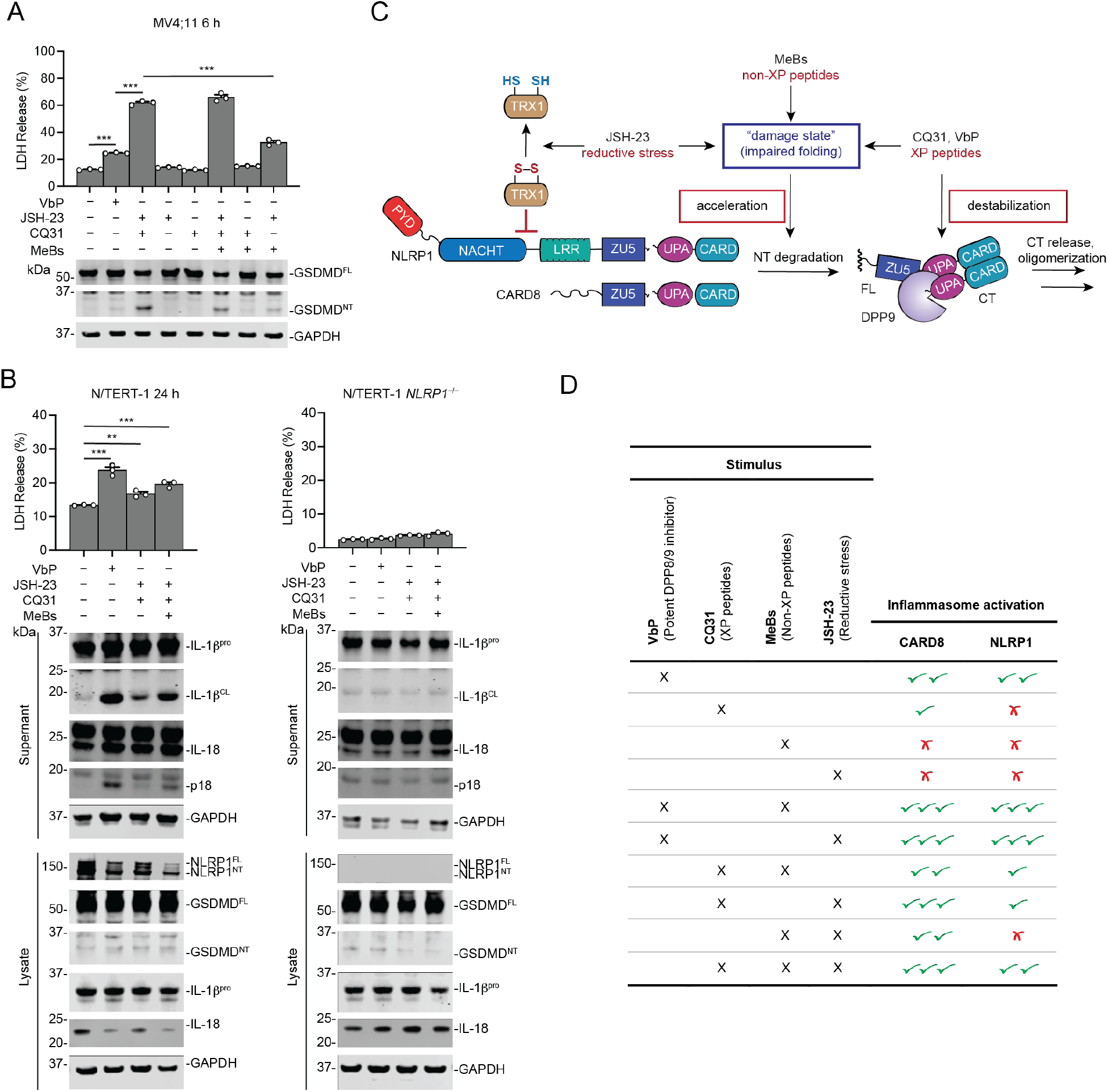
XP peptide accumulation and reductive stress activates NLRP1. (**A** and **B**) The indicated cells were treated with JSH-23 (2 μM), MeBs (10 μM), CQ31 (10 μM), (10 μM in **A**, 5 μM in **B**), or the specified combinations for the indicated time intervals. Data are means ± SEM of 3 replicates. *** *p* < 0.001, ** *p* < 0.01 by two-sided Students *t*-test. n.s., not significant. All data, including immunoblots, are representative of three or more independent experiments. (**C**) The proposed danger signals that CARD8 and NLRP1 detect. (**D**) Summary of small-molecule NLRP1 and CARD8 activators. See also Figure S7.

## DISCUSSION

We recently discovered that the oxidized, but not the reduced, form of TRX1 binds to and represses NLRP1 (Ball et al., 2021). As such, we predicted that reductive stress would abrogate this interaction and lead to inflammasome activation. However, no agents were known that induce reductive stress to test this hypothesis. Here, we identified and characterized several radical-trapping antioxidants, and in particular JSH-23, that induce reductive stress. To the best of our knowledge, these are the strongest known inducers of such reductive stress, and we predict they will gain widespread use to study this biological perturbation. Here, we used these probes to discover that reductive stress induces the degradation of NT fragments of both NLRP1 and CARD8, thereby releasing the CT fragments from autoinhibition. As CARD8 does not bind TRX1, these data indicate that reductive stress is monitored in at least two distinct ways, as discussed below.

The mechanisms that control the acceleration of NT degradation are not entirely clear. We recently proposed that MeBs (and likely VbP) induces the accumulation of proteasome-derived peptides, and that these peptides interfere with the folding of the NT domains, perhaps by inhibiting chaperones (Orth-He et al., 2022). Misfolded proteins are rapidly destroyed by the core 20S proteasome (Baugh et al., 2009; Liu et al., 2003), and therefore such misfolding would cause NT degradation. Consistent with this idea, we recently found that the core 20S proteasome, and not the ubiquitin-dependent 26S proteasome, controls CARD8 activation (Hsiao et al., 2022). We speculate that reductive stress similarly interferes with protein folding. Indeed, oxidative power is critical for protein folding in the endoplasmic reticulum (ER) (Tu and Weissman, 2004); it is possible that oxidative power plays a similar, but as yet unknown role in the folding of cytosolic proteins. Regardless, we predict that reductive stress impairs the folding of NLRP1 and CARD8’s NT fragments and sends them to the proteasome for destruction. In addition, we speculate that oxidized TRX1 stabilizes NLRP1 NT’s structure, thereby serving as a “checkpoint” to ensure misfolding is indeed associated with reductive stress. Future studies are needed not only to fully elucidate how reductive stress causes inflammasome activation, but also how it impacts proteostasis more generally.

Notably, our work here further confirms the importance of cytosolic ROS for the maintenance of homeostasis (Ray et al., 2012), but many mysteries regarding this ROS remain unanswered. For example, the identity, source, and regulation of the ROS that suppresses inflammasome activation are unknown. Furthermore, it is not yet clear what intracellular redox potential constitutes reductive stress, and how cells without inflammasomes respond to this perturbation. On that note, FNIP1 and CUL2^FEM1B^ were recently discovered as core components of the reductive stress response to mitochondrial inactivity (Manford et al., 2020). Projecting forward, it will be important to determine the relationships between cellular redox potential, FNIP1-CUL2^FEM1B^ signaling, TRX1 oxidation, and inflammasome activation.

Overall, our data here, coupled with our recent study on AP inhibitors, have now revealed that NLRP1 and CARD8 both detect at least two distinct danger signals: the accumulation of peptides and reductive stress. We speculate that these two signals are intimately related, and together contribute to a specific intracellular “damage state” in which proteins are unable to adopt and/or maintain folded structures. Moreover, we hypothesize NLRP1 interacts with DPP8/9 and TRX1 to verify that the damage state is indeed associated with those specific input signals. Lastly, we should note that, even though chemical probes have proved exceptionally useful in deconvoluting these danger signals, future studies are needed to understand how pathogens (and potentially other homeostasis-altering perturbations) induce reductive stress and peptide accumulation. Ultimately, we expect that these investigations will not only reveal key aspects of innate immunity, but will also uncover the mechanisms that regulate some of the most primordial processes in all of biology.

## Acknowledgements

This work was supported by the NIH (R01 AI137168, R01 AI163170, R01 CA266478 to D.A.B.; the MSKCC Core Grant P30 CA008748; T32 GM115327-Tan to E.L.O.-H), Gabrielle’s Angel Foundation (D.A.B.), Mr. William H. and Mrs. Alice Goodwin, the Commonwealth Foundation for Cancer Research, and The Center for Experimental Therapeutics of Memorial Sloan Kettering Cancer Center (D.A.B.), the Emerson Collective (D.A.B.).

## Author Contributions

D.A.B. conceived and directed the project. Q.W., J.C.H., N.Y., H.-C.H., C.M.O., E.L.O.-H., and D.P.B. performed cloning, gene editing, biochemistry, and cell biology experiments. Q.W., J.C.H., N.Y., and H.-C.H. designed experiments and analyzed data. D.A.B. and Q.W. wrote the manuscript. All authors reviewed and provided input to the manuscript.

## Materials and Methods

### Cell culture

HEK 293T, THP-1, and RAW 264.7 cells were purchased from ATCC. OCI-AML2 and MV4;11 cells were purchased from DSMZ. Naïve CD3 human T cells were purchased from HemaCare (Lot #21068415). N/TERT-1 cells were a gift from the Rheinwald Lab (Dickson et al., 2000). HEK 293T and RAW 264.7 cells were grown in Dulbecco’s Modified Eagle’s Medium (DMEM) with L-glutamine and 10% fetal bovine serum (FBS). Naïve human CD3 T cells, THP-1, MV4;11, and OCI-AML2 cells were grown in Roswell Park Memorial Institute (RPMI) medium 1640 with L-glutamine and 10% FBS. N/TERT-1 cells were grown in Keratinocyte serum-free medium (KSFM) supplemented with 1X penicillin/streptomycin, bovine pituitary extract (25 µg/mL) and epidermal growth factor (EGF) (0.2 ng/mL). All cells were grown at 37°C in a 5% CO_2_ atmosphere incubator. Cell lines were regularly tested for mycoplasma using the MycoAlert Mycoplasma Detection Kit (Lonza). *CARD8^−/−^*, *DPP8^−/−^*/*DPP9^−/−^*, and *CASP1^−/−^*THP-1 cells, *CASP1^−/−^* and *Nlrp1b^−/−^* RAW 264.7 cells, *CARD8^−/−^*, and *CASP1^−/−^* MV4;11, *NLRP1^−/−^* N/TERT-1 cells were generated as previously described (Ball et al., 2021; Johnson et al., 2018; Okondo et al., 2017). Doxycycline (DOX)-inducible CARD8 and NLRP1 WT and mutant knock-in *CARD8^−/−^* THP-1 were generated as previously described (Hollingsworth et al., 2021; Sharif et al., 2021).

### CRISPR/Cas9 gene editing

*DPP8/9*, *CARD8*, and *CASP1* knockout THP-1 cell lines and all HEK293T, RAW 264.7, N/TERT-1, and MV4;11 knockout cell lines were generated as previously described (Ball et al., 2021; Johnson et al., 2018; Okondo et al., 2017; Sharif et al., 2021). Briefly, 5 × 10^5^ HEK 293T cells stably expressing Cas9 were seeded in 6-well tissue culture dishes in 2 mL of media per well. The next day cells were transfected according to the manufacturer’s instructions (FuGENE HD, Promega) with 2 μg of the sgRNA plasmid(s). After 48 h, cells were transferred to a 10 cm tissue culture dish and selected with puromycin (1 μg/mL) until control cells were all dead. Single cell clones were isolated by serial dilution and confirmed by Western blot or sequencing, as indicated. To generate knockouts in RAW 264.7, MV4;11 and THP-1 cells, 1.5 × 10^6^ cells stably expressing Cas9 (Johnson et al., 2018) were infected with lentivirus containing sgRNA plasmids. After 48 h, cells were selected with puromycin (1 μg/mL) or hygromycin (100 μg/mL). Single cell clones were isolated by serial dilution and confirmed by Western blotting. *NLRP1* knockout N/TERT-1 keratinocytes were prepared by using the Neon Transfection System (ThermoFisher Scientific) following the manufacturer’s recommendations to deliver Cas9 ribonucleoprotein complexes containing an Alt-R CRISPR-Cas9 sgRNA and recombinant Cas9 (IDT). Briefly, sgRNA complexes were prepared by combining predesigned Alt-R CRISPR-Cas9 crRNA (NLRP1: 5’-CTGGATCCATGAATTGCCGG -3’) with Alt-R CRISPR-Cas9 tracrRNA to 44 µM and annealing by heating to 95 °C for 5 min followed by gradual cooling to ambient temperature over 30 min. To form the RNP complexes sgRNA samples and recombinant Alt-R Cas9 enzyme were combined and incubated for 20 min.

### Stable cell line generation

Cells stably expressing indicated protein constructs were generated by infection with lentivirus containing the desired plasmids. Briefly, the lentivirus was produced by transfecting 70% confluent HEK 293T cells with the desired plasmid along with psPAX2 and pMD2.G following the manufacturer’s instructions (FuGENE HD, Promega). The virus-containing medium was collected 48 h after transfection, passed through a 0.45 µm filter, and concentrated by PEG precipitation (Abcam). THP-1 cells were infected with the prepared lentivirus by centrifuging at 1000 × g for 1 h. After 48 h of incubation, cells were selected with an appropriate antibiotic.

### Transient transfections

HEK 293T cells were plated in 6-well tissue culture plates at 5.0 × 10^5^ cells/well in DMEM. The next day, the indicated plasmids were mixed with an empty vector to a total of 2.0 µg DNA in 125 µL Opti-MEM and transfected using FuGENE HD (Promega) according to the manufacturer’s protocol.

### Cloning

Plasmids for CARD8 WT and variants, CASP1, GSDMD, NLRP1 WT and variants, dTAG-CARD8^ZUC^ were cloned as described previously (Hollingsworth et al., 2021; Johnson et al., 2018; Sharif et al., 2021). Briefly, DNA sequences encoding the genes were purchased from GenScript, amplified by polymerase chain reaction (PCR), shuttled into the Gateway cloning system (ThermoFisher Scientific) using pDONR221 and pLEX307 vectors originating from pLEX307 (Addgene #41392). sgRNAs were designed using the Broad Institute’s web portal (Doench et al., 2016) (http://www.broadinstitute.org/rnai/public/analysis-tools/sgrna-design) and cloned into the lentiGuide-puro vector (Addgene #52963) as described previously (Sanjana et al., 2014). The sgRNA sequences used are described in the STAR Methods table.

### CellTiter-Glo and CytoTox-Fluor cytotoxicity assays

Cells were plated (2,000 cells per well) in white, 384-well clear-bottom plates (Corning) using an EL406 Microplate Washer/Dispenser (BioTek) in 25 μL final volume of the cell culture medium. Adherent cells were plated 12 h before treatment. To the cell plates were added compounds at different concentrations using a pintool (CyBio) and the plates were allowed to incubate for 1 h in the incubator before adding VbP (10 μM). After incubation for indicated times, CytoTox-Fluor reagent (Promega, G9262) was added according to the manufacturer’s protocol. The assay plates were then incubated for another 30 min before fluorescence was recorded using a Cytation 5 Cell Imaging Multi-Mode Reader (BioTek). Next, CellTiter-Glo reagent (Promega, G7573) was subsequently added to the assay plates following the manufacturer’s protocol. Assay plates were shaken on an orbital shaker for 2 min and incubated at 25 °C for 10 min. Luminescence was then read using a Cytation 5 Cell Imaging Multi-Mode Reader (BioTek).

### Excess over Bliss (EoB) calculation

EoB score is used to measure the combined effect of compound and VbP (Amzallag et al., 2019; Liu et al., 2018). EoB ˃ 0 indicates the compound synergizes with VbP-induced cell death, EoB < 0 indicates the compound blocks VbP-induced cell death, and it was calculated using the following equations:

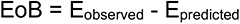

Where E_observed_ is the observed cell death for the treatment of compound + VbP and was calculated by subtracting the observed viability based on CTG assay from 100. E_predicted_ was calculated based on the following equation:

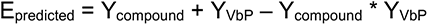

where Y_compound_ is the observed cell death with compound alone at indicated dose and Y_VbP_ is the observed cell death with VbP (10 μM) alone. Y_compound_ and Y_VbP_ are calculated by subtracting the observed viability for compound and VbP based on CTG assay from 100, respectively. All compounds were tested in triplicate at indicated doses.

### Propidium iodide uptake analysis

2 × 10^4^ MV4;11 or THP-1 cells were plated in 384-well, black, clear-bottom plates (Corning) in 40 μL of RPMI medium. RAW 264.7 cells were plated 12 h before treatment. Cells were then treated as indicated and 40 μL propidium iodide (PI) at 20 μM was added to each well. PI fluorescence was measured at Ex/Em: 535/617 nm at 37°C every 5 min using Cytation 5 Cell Imaging Multi-Mode Reader (BioTek). The obtained measurements were baseline corrected to DMSO at each timepoint and was presented as F_treatment_ - F_DMSO_. F_treatment_, Fluorescence measurement for compound treatment at time t; F_DMSO_, fluorescence measurement for DMSO control at time t.

### LDH cytotoxicity assays

MV4;11, THP-1, OCI-AML2, RAW 264.7, and RAW 264.7^ASC^ cells were plated in 12-well tissue culture plates at 5 × 10^5^ cells/well, Naïve human CD3 T cells were plated in 12-well tissue culture plates at 2 × 10^6^ cells/well. N/TERT-1 keratinocytes were seeded at 2 × 10^5^ cells/well (in 2 mL medium) in 6-well tissue culture plates. RAW 264.7, and RAW 264.7^ASC^ cells were incubated overnight before treatment. N/TERT-1 keratinocytes were incubated 48h before treatment. *CARD8^−/−^* THP-1 cells containing a DOX-inducible CARD8 WT, NLRP1 WT, CARD8 E274R, or NLRP1 P1214R protein were seeded at 2.5 × 10^5^ cells/well in 12-well tissue culture dishes. The cells were then pre-incubated with the indicated concentration of doxycycline for indicated times before compound treatment. For all cells, after treatment with indicated compounds for indicated times, supernatants were analyzed for LDH activity using the Pierce LDH Cytotoxicity Assay Kit (Life Technologies). LDH activity was quantified relative to a lysis control where cells were lysed by adding 8 µL of a 9% Triton X-100 solution.

### Ferroptosis assays

RAW 264.7 cells were plated (2,000 cells per well) in white, 384-well clear-bottom plates (Corning) manually in 25 μL final volume of the cell culture medium. After overnight incubation, cell plates were allowed to pre-treat with compounds using a pintool (CyBio) for 30 min in the incubator before adding RSL3 (0.25 µM). After 4.5 h, cytotoxicity was assessed by CTF and CTG assay as mentioned in the CellTiter-Glo and CytoTox-Fluor cytotoxicity assays part.

### DCFDA/H_2_DCFDA cellular ROS assay

This assay was carried out using a cellular ROS kit (Abcam, Ab113851) according to the manufacturer’s instructions. Briefly, 1 X ROS assay buffer was freshly prepared and DCFDA solution was diluted to 20 μM with this newly prepared buffer. *CASP1^−/−^* MV4;11 or THP-1 cells were collected, washed with PBS, suspended in 20 μM DCFDA solution (1 mL per million cells), and incubated for 30 min. Cells were then centrifuged, washed with PBS twice, suspended in corresponding cell culture medium (supplemented with 10% FBS) without phenol red, and plated (10,000 cells/well) in black, 384-well clear-bottom plates (Corning) manually in 25 μL final volume before treated with compounds. Fluorescence was measured at Ex/Em: 485/535 nm recorded at 37°C every 5 min using Cytation 5 Cell Imaging Multi-Mode Reader (BioTek).

### Cell-free 2,2-diphenyl-1-picrylhydrazyl (DPPH) assay

Freshly prepared DPPH solution in methanol (40 μM) was plated on white, 384-well clear-bottom plates. Methanol was utilized as a background control. Compounds were added at different concentrations using a pintool (CyBio) and the cell plate was sat in dark for 1 h before absorbance at 517 nm was recorded using Cytation 5 Cell Imaging Multi-Mode Reader (BioTek). All values were normalized to the background. Each condition was tested in triplicate.

### C11 BODIPY 581/591 assay

*CASP1^−/−^* MV4;11 cells were seeded at 1 × 10^6^ in a 6-well plate and treated for 2.5 h with the indicated compounds before staining with 5 μM of C11 BODIPY 581/591 for an additional 30 min. Cells were subsequently collected, washed twice with FACS buffer (PBS with 2.5% FBS), and analyzed for the FITC and PE-CF594 fluorochromes using the Fortessa (BD Biosciences). Unstained, DAPI only, and C11 BODIPY 581/591 only technical controls were included to ensure proper gating. The gating pipeline starts by isolating the bulk cell population and excluding doublets, then choosing the population that was DAPI negative before plotting the histogram of cells corresponding to the FITC fluorochrome. To note, the C11-BODIPY dye emits a green wavelength when oxidized.

### AMC substrate assays

For recombinant enzyme assays, 25 μL DPP9 enzyme solution (1.0 nM) was added to a 384-well, black, clear-bottom plate (Corning). Compounds at different concentrations were added using a pintool (CyBio) and the enzyme plate was shaken at room temperature for 45 min, followed by Ala-Pro-AMC (25 μM) substrate to initiate the reaction. For in cell assays, 8.0 × 10^4^ *CASP1*^-/-^ MV4;11 or HEK 293T cells were seeded per well in a 96-well, black, clear-bottom plate (Corning) (overnight for HEK 293T cells) in OptiMEM and treated with compounds for 5 h, and then with Sitagliptin (1 μM) to block DPP4 activity for 1h before substrate (Ala-Pro-AMC, 5 μM; Ala-AMC, 100 μM) was added to the media to initiate the reaction. AMC fluorescence (Ex/Em: 380/460 nm) was recorded at 25 °C for 30 - 60 mins. Cleavage rates are reported as the slope of the linear regression of AMC fluorescence vs. time data curve.

### IL-1β ELISA assays

2 × 10^5^ N/TERT-1 keratinocytes in 2 mL cell culture medium were plated on 6-well tissue culture plates and incubated for 48 h. Cells were then treated with compounds for the indicated time points before LDH release analysis. 200 μL spent media sample were collected, centrifuged at 1000 g for 1 min, and the supernatants were utilized for IL-1β quantification using the R&D Human IL-1β quantikine ELISA kit according to the manufacturer’s instructions.

### Immunoblotting

Cells were washed 2 × in PBS (pH = 7.4), resuspended in PBS, and lysed by sonication. Protein concentrations were determined and normalized using the DCA Protein Assay kit (Bio-Rad). To prepare supernatant western blotting samples of N/TERT-1 keratinocytes, spent media of each treatment were combined, centrifuged at 400 g for 3 min, and the supernatant was precipitated by addition of four sample volumes of acetone at -20 °C overnight followed by centrifugation at 3000 g for 30 min at 4 °C and decanting. Protein pellets were then suspended in 1X PBS and combined 1:1 with 2X sample loading buffer before heating to 97 °C for 10 min.

Samples were run on NuPAGE 4 to 12%, Bis-Tris, 1.0 mM, Midi Protein Gel (Invitrogen) for 45 - 60 min at 175 V. Gels were transferred to nitrocellulose with the Trans-Blot Turbo Transfer System (BIO-RAD). Membranes were blocked with Intercept (TBS) Blocking Buffer (LI-COR) for 30 min at ambient temperature, before incubating with primary antibody overnight at 4°C. Blots were washed 3 times with TBST buffer before incubating with secondary antibody for 60 min at ambient temperature. Blots were washed 3 times, rinsed with water, and imaged via Odyssey CLx (LI-COR).

### dTAG-CARD8 Assay

HEK 293T cells stably expressing CASP1 and GSDMD were seeded at 1.5 × 10^5^ cells per well in 12-well tissue culture dishes. After 48 h, the cells were transfected with plasmids encoding dTAG-CARD8-ZUC (0.5 μg), CARD8 FIIND-S297A (0.3 μg), and RFP (0.2 μg) with FuGENE HD, according to the manufacturer’s instructions (Promega). After 24 h, cells were treated with DMSO, dTAG13 (500 nM), and indicated compounds for 3 h prior to LDH release and immunoblot analyses.

### Statistical analysis

Two-sided Student’s t tests were used for significance testing unless stated otherwise. *P* values less than 0.05 were considered to be significant. Graphs and error bars represent means ± SEM of three independent experiments unless stated otherwise. The investigators were not blinded in all experiments. All statistical analysis was performed using Microsoft Excel and GraphPad Prism 9.

