## Supplemental Figures 1-7 for "The NLRP1 and CARD8 inflammasomes detect reductive stress"

**SUPPLEMENTARY INFORMATION**

**This file contains:**
**Figure S1-S7**

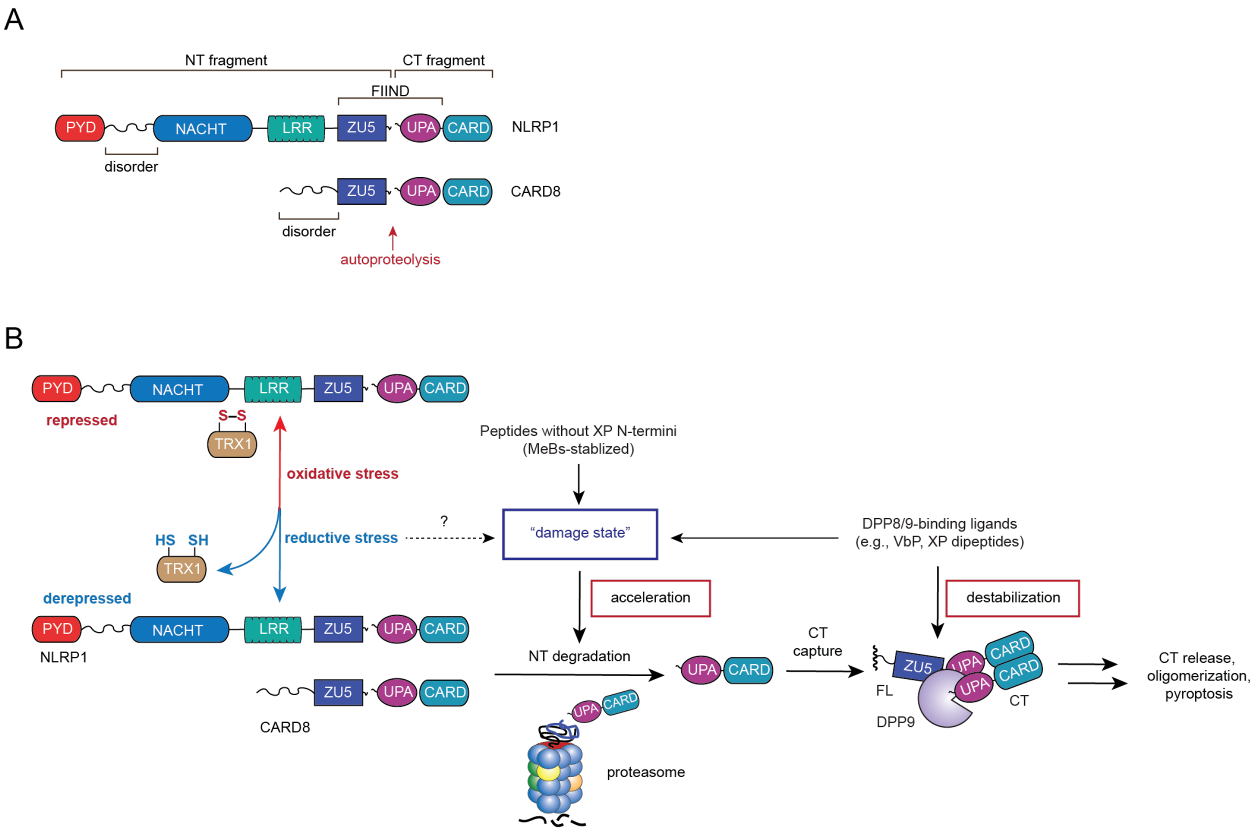


**Figure S1. Overview of NLRP1 and CARD8 activation.** (**A**) Domain organizations of human NLRP1 and CARD8. The autoproteolysis sites are indicated. (**B**) The accelerated degradation of the NT fragments and the destabilization of the DPP8/9 ternary complexes activates the NLRP1 and CARD8 inflammasomes. We propose that at least two distinct signals – peptide accumulation and reductive stress – create a damaged state that accelerates NT degradation. XP peptides also destabilize the DPP8/9 ternary complex.

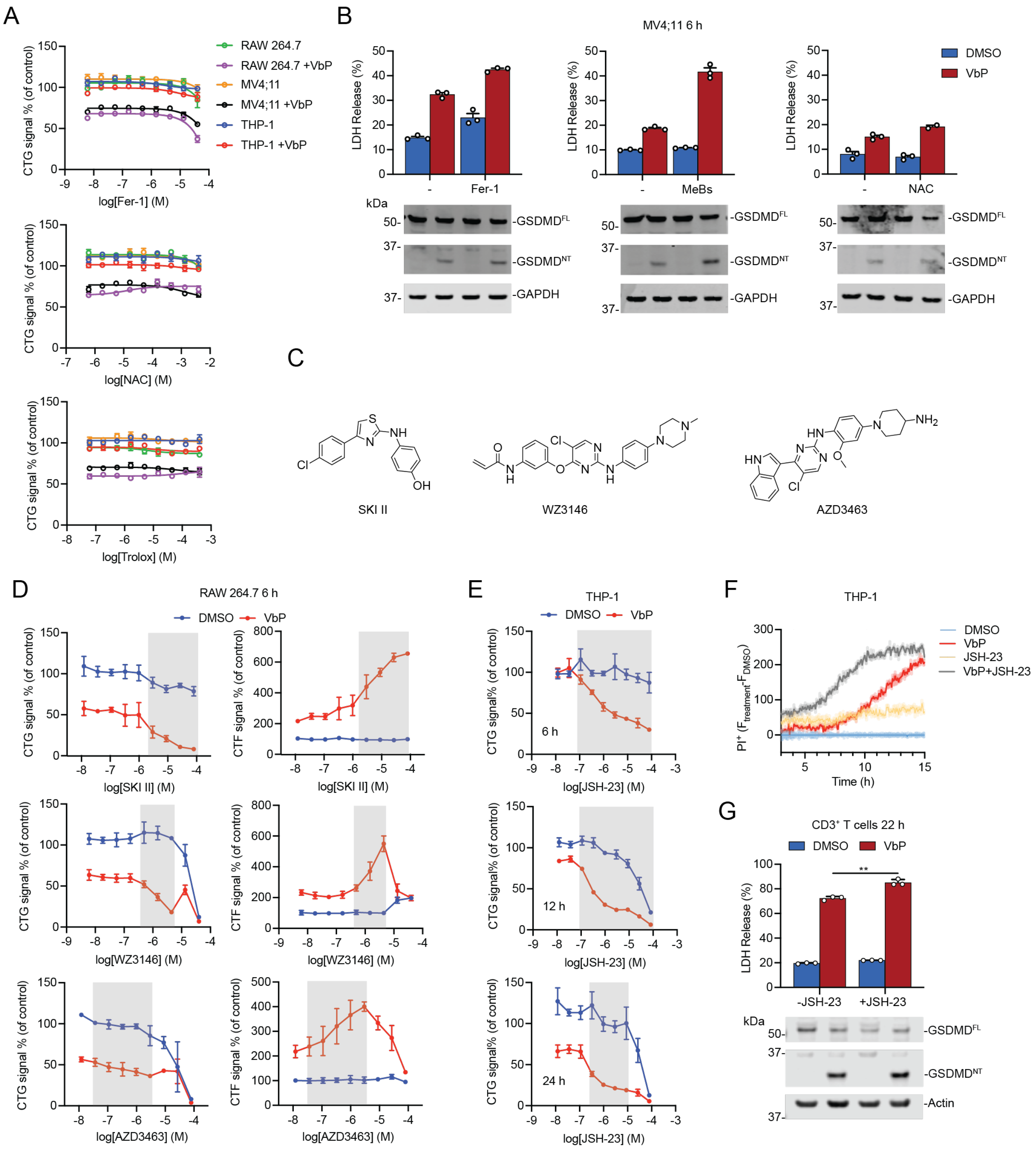


**Figure S2. Ferroptosis inhibitors enhance VbP-induced cell death, related to Figure 1.** (**A** and **B**) The indicated cells were treated with Fer-1, NAC, MeBs, and Trolox ± VbP (10 μM). Cell death was evaluated after 6 h by CTG (**A**) and by LDH release and immunoblot analyses (**B**). In **B**, cells were pre-incubated with Fer-1 (40 μM), MeBs (20 μM), and NAC (2 mM) for 1 h before the addition of VbP (10 μM). (**C**) Structures of SKI II, WZ3146, and AZD3463. (**D** and **E**) The indicated cells were treated with SKI II, WZ3146, and AZD3463 ± VbP (10 μM) for 6 h before cell death was assessed by CTG and CTF. Synergistic cell death is highlighted in gray. (**F**) THP-1 cells were treated with JSH-23 (10 μM), VbP (10 μM), or both for 3 h before PI uptake was monitored for 12 h. (**G**) Resting human $CD3^+$ T cells were treated with JSH-23 (10 μM), VbP (0.2 μM), or both for 22 h before LDH release and immunoblot analyses. Data are means ± SEM of 3 replicates. ** $p < 0.01$ by two-sided Students *t*-test. All data, including immunoblots, are representative of three or more independent experiments.

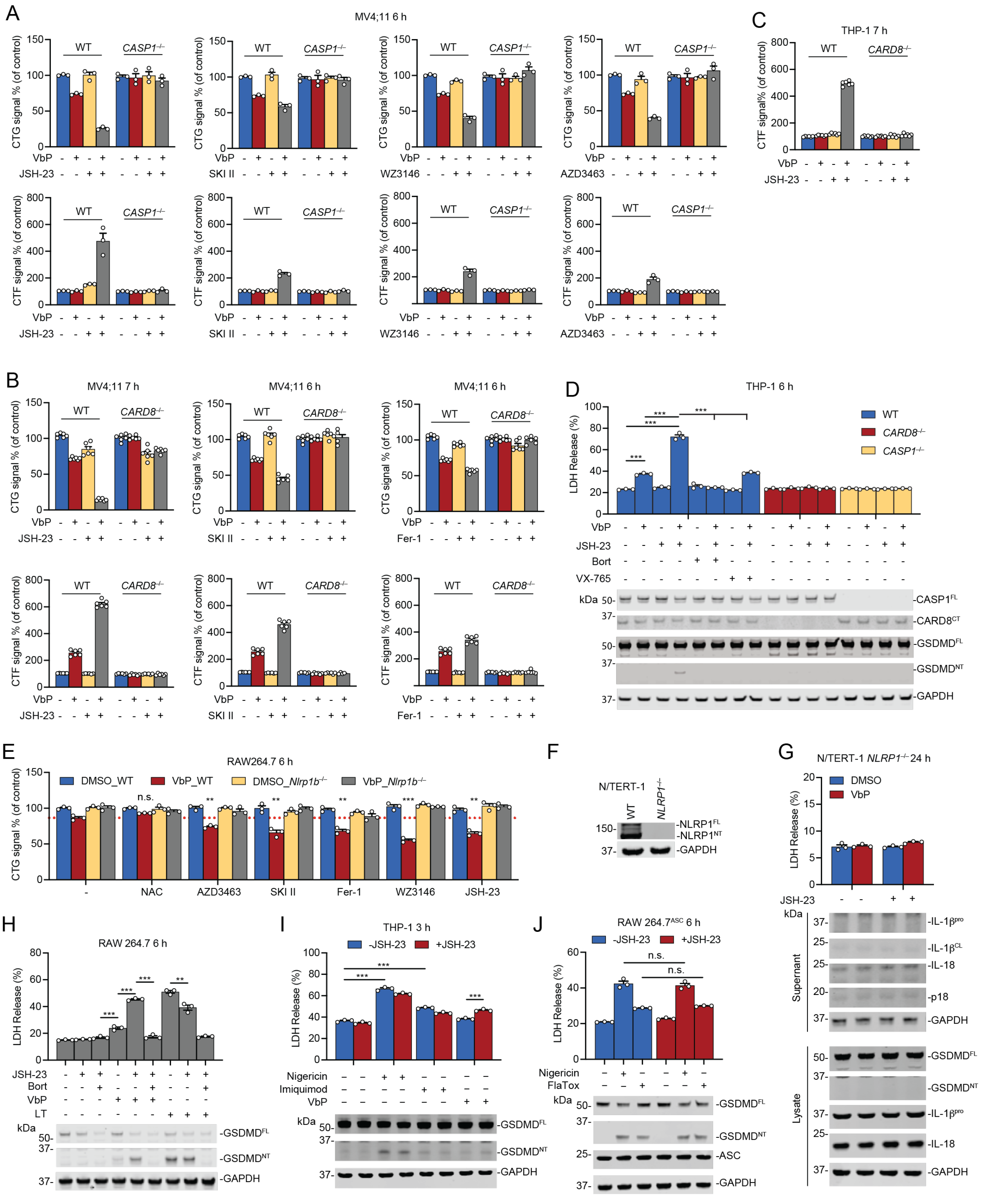

**Figure S3. Ferroptosis inhibitors selectively synergize with VbP, related to Figure 2.** (**A-E**) The indicated cell types were treated with VbP (10 μM), JSH-23 (10 μM), SKI II (10 μM in **A** and **B**; 20 μM in **E**), WZ3146 (4 μM), AZD3463 (1 μM), Fer-1 (20 μM in **B**; 40 μM in **E**), NAC (4 mM), or the specified combinations for the indicated time intervals prior to CTG, CTF, LDH release, and immunoblot analyses. In **D**, bortezomib (Bort, 1 μM) and VX-765 (10 μM) were added 15 min prior to VbP and JSH-23. The red dashed line in **E** indicates VbP-induced death. (**F**) Confirmation of *NLRP1* knockout by immunoblotting. (**G**) *NLRP1*$^{-/-}$ N/TERT-1 were treated with JSH-23 (2 μM), VbP (0.1 μM), or both for 24 h before LDH release and immunoblot analyses. (**H**) RAW 264.7 cells were treated with JSH-23 (2.5 μM), VbP (10 μM), or both for 5 h. Alternatively, cells were treated with anthrax lethal toxin (LT, 1 μg/mL) in the presence or absence of JSH-23 (2.5 μM) for 3 h. Bort (20 μM) was added to the indicated samples 15 min prior to VbP and 2.5 h prior to LT. Cell death was evaluated by LDH release and immunoblot analyses. (**I**) THP-1 cells were primed with LPS (1 μg/mL, 14 h) and then treated with DMSO, nigericin (1 μM), or imiquimod (30 μM) ± JSH-23 (2 μM) for 3 h prior to LDH release and immunoblot analyses. (**J**) RAW 264.7 cells stably expressing ASC were primed with LPS (1 μg/mL, 14 h) and then treated with DMSO, nigericin (10 μM), or LFn-flagellin ("FlaTox", 1 μg/mL) ± JSH-23 (2 μM) for 6 h before LDH release and immunoblot analyses. Data are means ± SEM of 3 replicates. *** $p < 0.001$, ** $p < 0.01$ by two-sided Students *t*-test. n.s., not significant. All data, including immunoblots, are representative of three or more independent experiments.

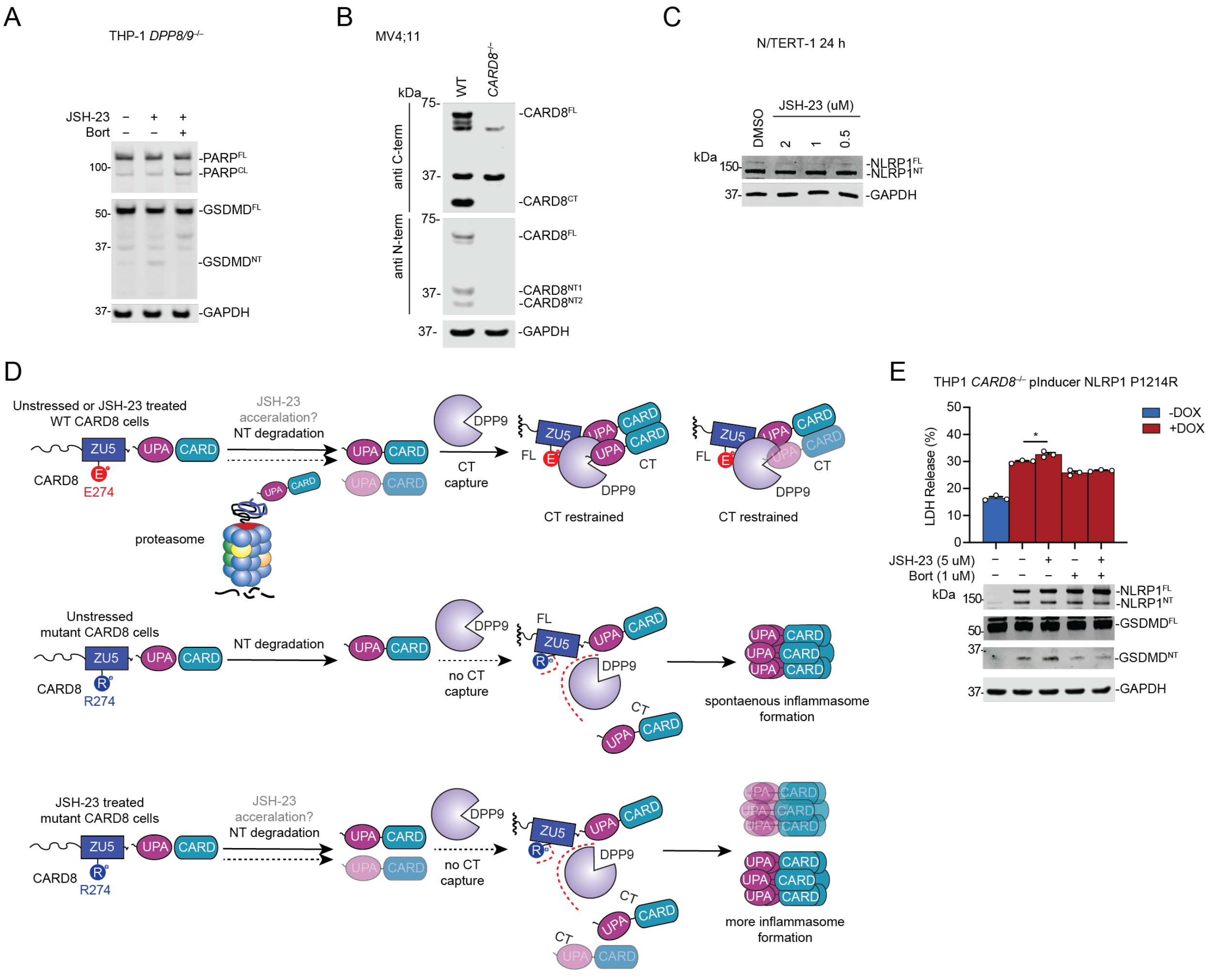


**Figure S4. JSH-23 promotes CARD8 and NLRP1 NT degradation, related to Figure 3.**

(**A**) *DPP8/9*$^{-/-}$ THP-1 cells were treated with JSH-23 (2.5 μM) ± Bort (1 μM) for 6 h before LDH release and immunoblot analyses. (B) Confirmation of *CARD8* knockout in MV4;11 cells by immunoblotting. (**C**) N/TERT-1 keratinocytes were treated with JSH-23 at the indicated concentrations for 24 h protein levels were assessed by immunoblotting. (**D**) Schematic of DOX-inducible CARD8 E274R assay used to evaluate accelerated CARD8 NT degradation. (**E**) *CARD8*$^{-/-}$ THP-1 cells containing a DPP9 non-binding NLRP1 P1214 mutant were treated with or without DOX (1 ug/mL, 2 h) followed by Bort (1 μM), JSH-23 (5 μM), or both for 6 h prior to LDH release and immunoblot analyses. Data are means ± SEM of 3 replicates. * $p < 0.05$ by two-sided

Students *t*-test. n.s., not significant.  All data, including immunoblots, are representative of three or more independent experiments.

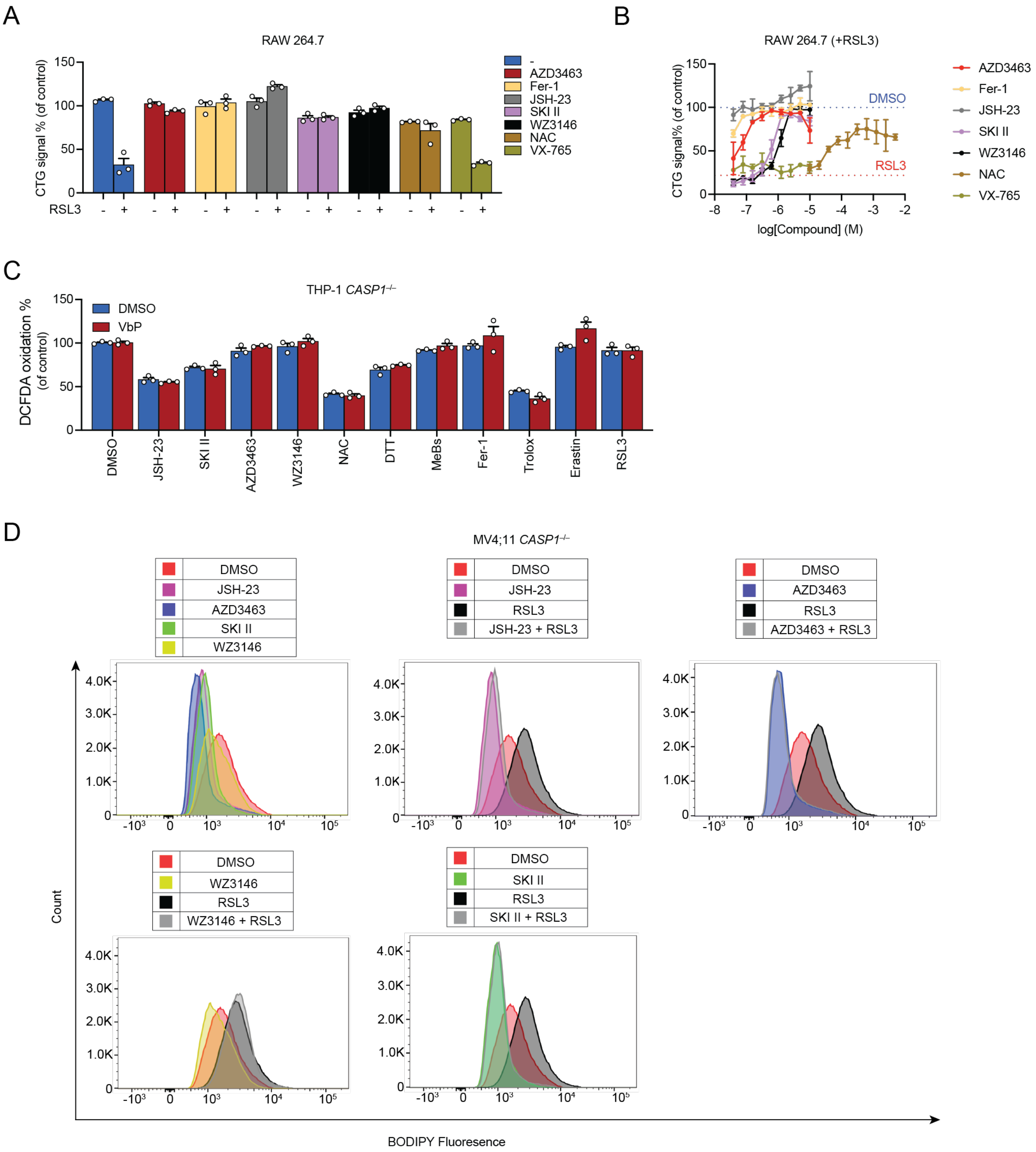


**Figure S5. Synergistic compounds reduce cytosolic and lipid ROS, related to Figure 4.** (**A** and **B**) RAW 264.7 cells were treated with the indicated compounds ± RSL3 (0.25 µM). Cell death was evaluated by CTG after 4.5 h. All compounds in **A** were tested at 5 µM (except for NAC at 1.2 mM). (**C**) VbP (10 µM), JSH-23 (4 µM), SKI II (20 µM), AZD3463 (4 µM), WZ3146 (0.8 µM),

NAC (1 mM), DTT (1 mM), MeBs (20 μM), Fer-1 (0.8 μM), Trolox (400 μM), Erastin (20 μM), and RSL3 (0.5 μM) ± VbP (10 μM) were tested for their impact on the cell-permeable $H_2$DCFDA probe in *CASP1*$^{-/-}$ THP-1 cells. (**D**) The indicated compounds (RSL3 was tested at 0.5 μM, others were tested at 1 μM) were tested for their impact on lipid ROS at 3 h in *CASP1*$^{-/-}$ MV4;11 using the C11 BODIPY 581/591 probe. Data are means ± SEM of 3 replicates. All data are representative of three or more independent experiments.

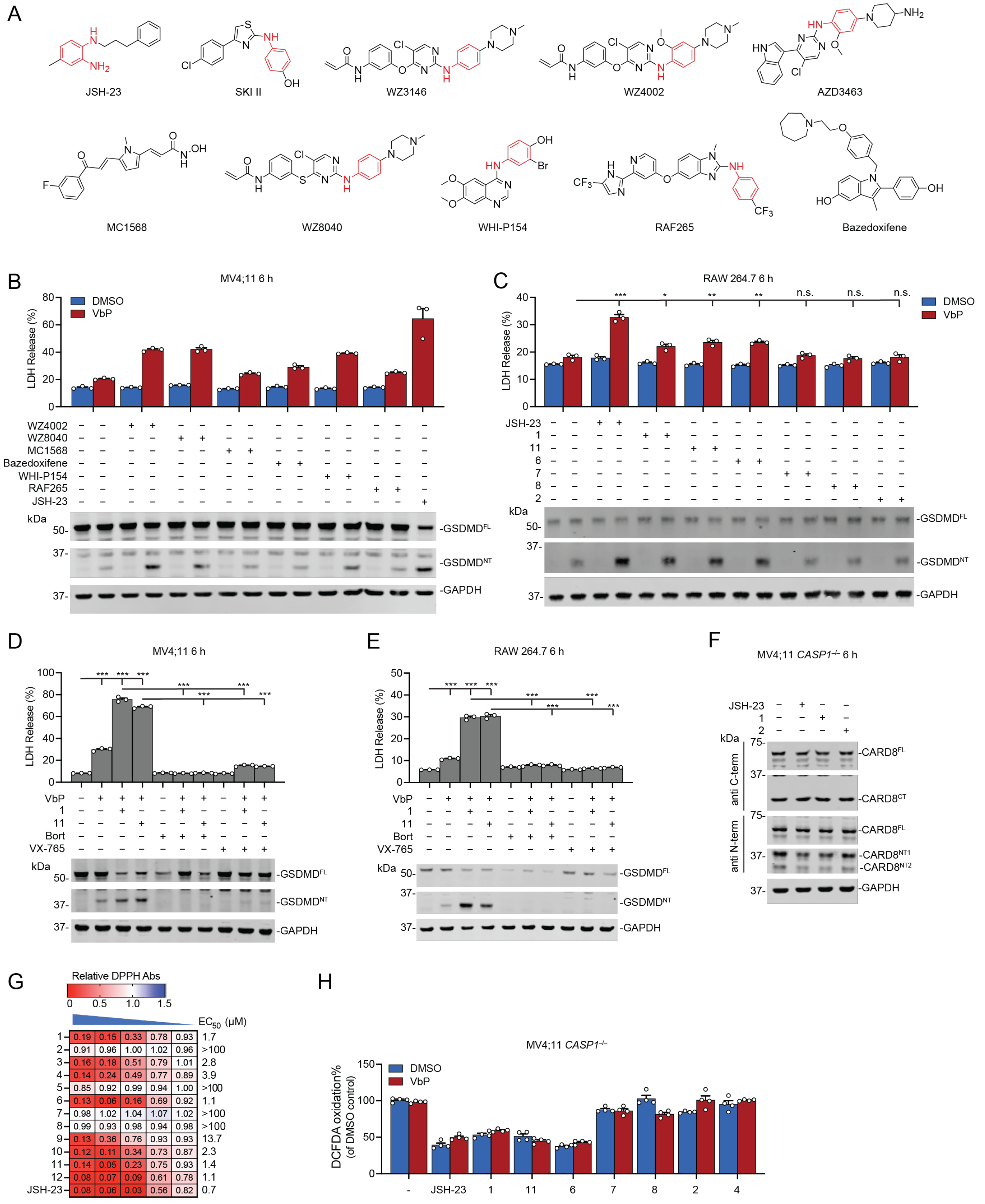


**Figure S6. Conjugated amines are critical for RTA activity, related to Figure 5.** (**A**) Structures of JSH-23 and other synergistic or non-synergistic compounds from **Figure 1D**. Many of the synergistic compounds have conjugated secondary amines (colored red). (**B**) MV4;11 cells were treated with WZ4002 (3 μM), WZ8040 (1.5 μM), MC1568 (3 μM), bazedoxifene (4.5 μM), WHI-P154 (1 μM), RAF265 (4.5 μM), or JSH-23 (2 μM) ± VbP (10 μM) for 6 h prior to LDH release and immunoblot analyses. (**C**) RAW 264.7 cells were treated with the indicated compounds (all at 5 μM) ± VbP (10 μM) for 6 h prior to LDH release and immunoblot analyses. (**D** and **E**) MV4;11 (**D**) or RAW 264.7 (**E**) cells were pre-treated with Bortezomib (Bort, 1 μM) or VX-765 (25 μM) for 30 min before applying VbP (10 μM), **1** (5 μM), **11** (5 μM), or the specified combinations for 6 h. Pyroptosis was assessed by LDH release and immunoblot analyses. (**F**) *CASP1*$^{-/-}$ MV4;11 cells were treated with JSH-23, **1**, or **2** (all at 2 μM) for 6 h. CARD8 protein levels were determined by immunoblotting. CARD8 NT1 and NT2 are from different splice isoforms of CARD8. (**G**) Compounds were tested (all at 40 μM in 3-fold dilution series) for radical trapping activity using a cell-free DPPH assay. The $EC_{50}$ values are shown. (**H**) Compounds were tested (all at 5 μM) ± VbP (10 μM) for their impact on the oxidation of the cell-permeable $H_2DCFDA$ probe in *CASP1*$^{-/-}$ MV4;11 cells. Data are means ± SEM of 3 replicates. *** $p < 0.001$, ** $p < 0.01$, * $p < 0.05$ by two-sided Students *t*-test. n.s., not significant. All data, including immunoblots, are representative of three or more independent experiments.

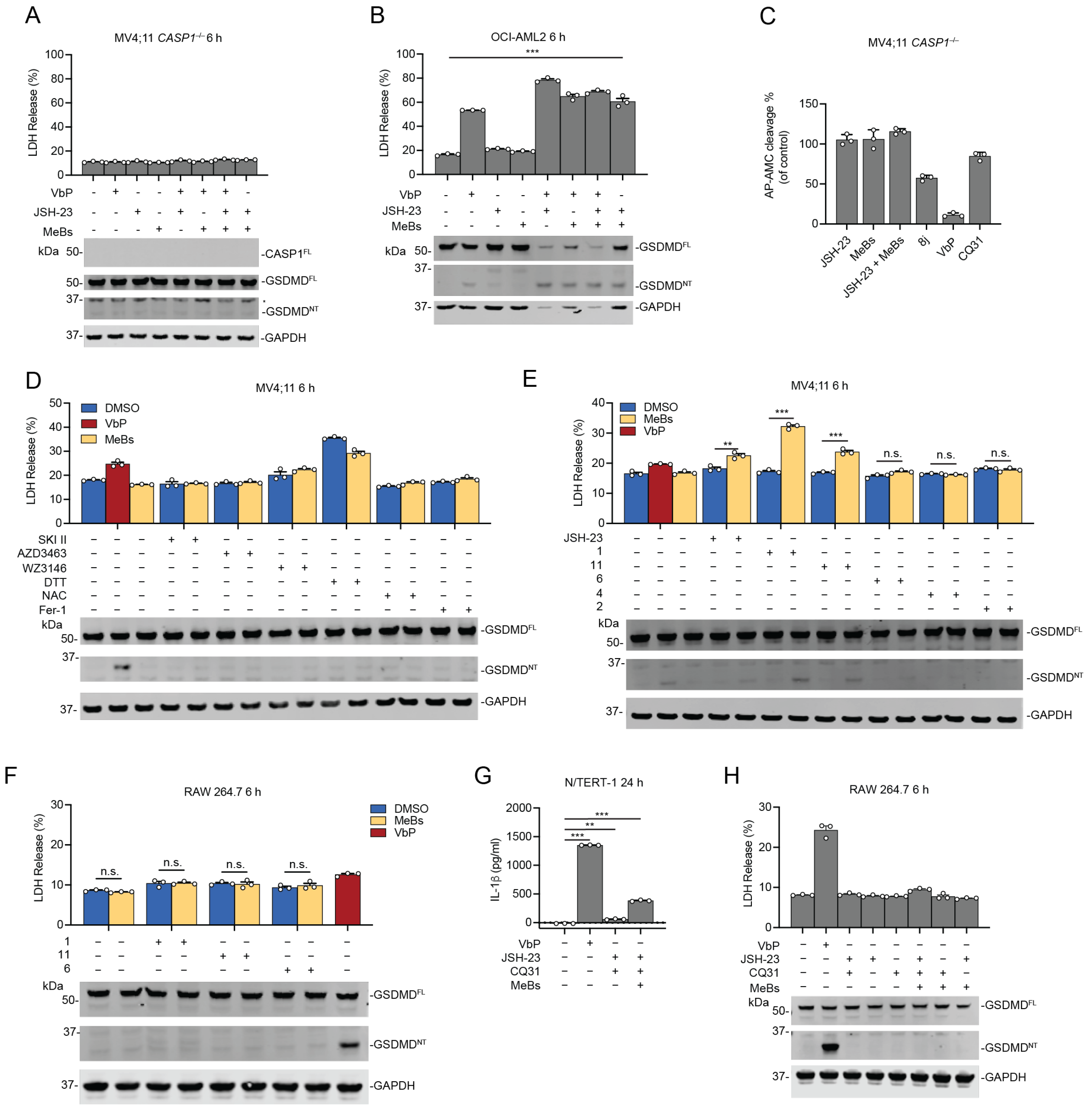

**Figure S7. AP inhibitors and RTAs specifically activate CARD8, not NLRP1, related to Figure 6 and 7.** (**A** and **B**) The indicated cells were treated with JSH-23 (2 μM), MeBs (10 μM), VbP (10 μM), or the specified combination for 6 h before LDH release and immunoblot analyses. (**C**) JSH-23 (2 μM) and MeBs (10 μM), unlike 8j (10 μM), VbP (10 μM) and CQ31 (10 μM), do not inhibit the activity of DPP8/9 in *CASP1*$^{-/-}$ MV4;11 cells as measured by AP-AMC cleavage. (**D-F**) SKI II (20 μM), AZD3463 (1 μM), WZ3146 (5 μM), DTT (2 mM), NAC (2 mM), Fer-1 (20 μM), JSH-

23 (2 μM), **1** (15 μM), **11** (15 μM), **6** (15 μM), **4** (15 μM), and **2** (15 μM) were added to the indicated cells alone or in combination with MeBs (10 μM). VbP (10 μM) was used as a control. Cell death was determined after 6 h using LDH and immunoblot analyses. (**G** and **H**) The indicated cells were treated with VbP (5 μM in **G**, 10 μM in **H**), JSH-23 (2 μM), MeBs (10 μM), CQ31 (10 μM) alone or in the specified combinations for the indicated time intervals before assessing pyroptosis using IL-1β (**G** or LDH release and immunoblot analyses (**H**). Data are means ± SEM of 3 replicates. *** $p < 0.001$, ** $p < 0.01$ by two-sided Students *t*-test. n.s., not significant. All data, including immunoblots, are representative of three or more independent experiments.
